# Error-free and efficient prime editing in absence of maternal Polθ

**DOI:** 10.64898/2026.02.02.703001

**Authors:** François Kroll, Malo Serafini, Luna de Barbarin, José Roberto Alvarez Vargas, Julie Thanh-Mai Dang, Marion Rosello, Marie As, Anne De Cian, Jean-Paul Concordet, Carine Giovannangeli, Filippo Del Bene

## Abstract

Prime editing is a versatile genome-editing technology developed to make small edits while minimising the unwanted mutations associated with approaches based on double-strand breaks. In zebrafish embryos, however, prime editing is highly error-prone, typically yielding more unwanted mutations than precise edits. Here, we show that most unwanted mutations are deletions flanked by microhomologies and insertions templated from neighbouring sequences, both hallmarks of microhomology-mediated end-joining of double-strand breaks. These breaks are unlikely to result from direct cleavage of both genomic strands by the prime editor. Instead, we propose that the rapid cell divisions of early zebrafish embryos promote the conversion of nicks into double-strand breaks by replication forks. Loss of maternally deposited DNA polymerase θ (Polθ), the core factor in microhomology-mediated end-joining, abolishes unwanted mutations and can increase edit rates above 50%, resulting in error-free and efficient prime editing. Our work identifies the source of unwanted prime-editing mutations in zebrafish and establishes a practical strategy to achieve high edit rates and near-perfect precision.

## Introduction

Owing to their rapid external development, zebrafish larvae have become widely used to model genetic diseases in biomedical research. Generation of loss-of-function “knockout” alleles using standard CRISPR/Cas9 in zebrafish is now highly efficient (>90% mutation rate for most targets), even enabling the direct use of injected animals in experiments ^1,2^. However, only ∼8% of coding human variants lead to a premature stop codon (Supplementary Fig. 1). In most cases, the exact variant observed in the patient should thus be introduced in the zebrafish genome to model the disease or confirm the variant’s pathogenicity, necessitating precise editing. To do so, researchers have extensively employed homology-directed repair (HDR) of a Cas9-mediated double-strand break (DSB). Although the approach has been under development for a decade ^3,4^, it still achieves <10% edit rate in most cases ^5,6^. Because zebrafish embryos repair most DSBs using Polθ-dependent microhomology-mediated end-joining (MMEJ), rather than HDR or non-homologous end-joining (NHEJ) ^7,8^, the majority of induced mutations are unwanted insertions and deletions (indels), most of which (∼66%) cause frameshifts leading to premature stop codons.

To avoid using DSBs, genome editing techniques based on single-strand breaks—nicks—have been developed, namely base editing and prime editing. Base editing can achieve high rates of precise edits in zebrafish ^9,10^ but regularly cause “bystander edits” and can only mediate 4 of the 12 possible single-nucleotide substitutions; other substitutions or small insertions and deletions are not feasible. For small edits, a potentially universal approach is prime editing ^11^. Prime editing fuses a Cas9 nickase to a reverse transcriptase (RT). The guide RNA (gRNA), now called prime-editing guide RNA (pegRNA), is extended to include the primer binding site (PBS) and the template for the RT (RTT) encoding the desired edit. Cas9 nicks the genome at the locus specified by the pegRNA’s spacer, releasing a 3′ end that anneals to the PBS, allowing the RT to polymerise the end using the RTT as template, thereby writing the desired edit into one genomic strand. The edited 3′ flap must then anneal to the other strand by replacing the unedited 5′ flap, which is excised by an endogenous flap endonuclease. Ligation of the edited flap leads to a heteroduplex. The edit is eventually incorporated in the other strand by DNA mismatch repair or at the next cell cycle when the edited strand is used as template strand. Previous work has shown the potential of prime editing in zebrafish ^5,6,12–14^. However, reported edit rates are typically low, particularly so for substitutions (<5% per embryo), which is the largest class of human variants (>90%, Supplementary Fig. 1). In contrast to human cells, rates of unwanted indels are surprisingly high in zebrafish embryos, often several times the rate of the desired edit, undermining the main promise of genome editing without DSBs.

Here, we report that virtually every indel induced by prime editing in zebrafish was dependent on Polθ (Polq) activity during MMEJ. Prime editing in *polq* mutants can achieve edit rates above 50% and is almost perfectly precise; that is, the only observed mutation is the desired edit.

## Results

### Optimising zebrafish prime editing using a pigmentation-based assay

We first aimed to improve prime editing rates in zebrafish. We previously generated an *slc45a2* knockout line by mutating a tryptophan codon (TGG, W121) into a stop (TAA, *) ^9^. The resulting *slc45a2^W121*^* line recapitulated the *albino* phenotype caused by reduced melanin production ^15^. Inspired by work on HDR ^3^, we reasoned that using prime editing to restore the wild-type allele (Fig. 1a) in homozygous *slc45a2^W121*^* embryos would make pigmentation reappear in patches, providing a rapid phenotypic readout of precise (pure) edits. Most unwanted indels are expected to leave cells unpigmented as they do not restore Slc45a2’s function. To validate this “pigmentation recovery assay”, we injected PE2 protein complexed with a synthetic pegRNA encoding the AA>GG edit in homozygous *slc45a2^W121*^* embryos. At 3 days post-fertilisation (dpf), we scored eye pigmentation (Fig. 1b) and sequenced individual larvae assigned to each score (Supplementary Fig. 2a). We classified reads as reference, pure edit, impure edit, or scaffold incorporation if we detected pegRNA scaffold-templated nucleotides (Supplementary Fig. 2b). The scaffold-templated nucleotides could appear as insertions or substitutions, which we commonly refer to as scaffold incorporations (Supplementary Fig. 2c). As intended, increasing pigmentation scores correlated with higher pure edit rates (1.4–8.8%, Fig. 1c) but not with indel rates (Fig. 1d). Scores were consistent between experimenters (Supplementary Fig. 2d). The pigmentation recovery assay can therefore be used to quickly estimate pure prime editing rates.

**Figure 1.**
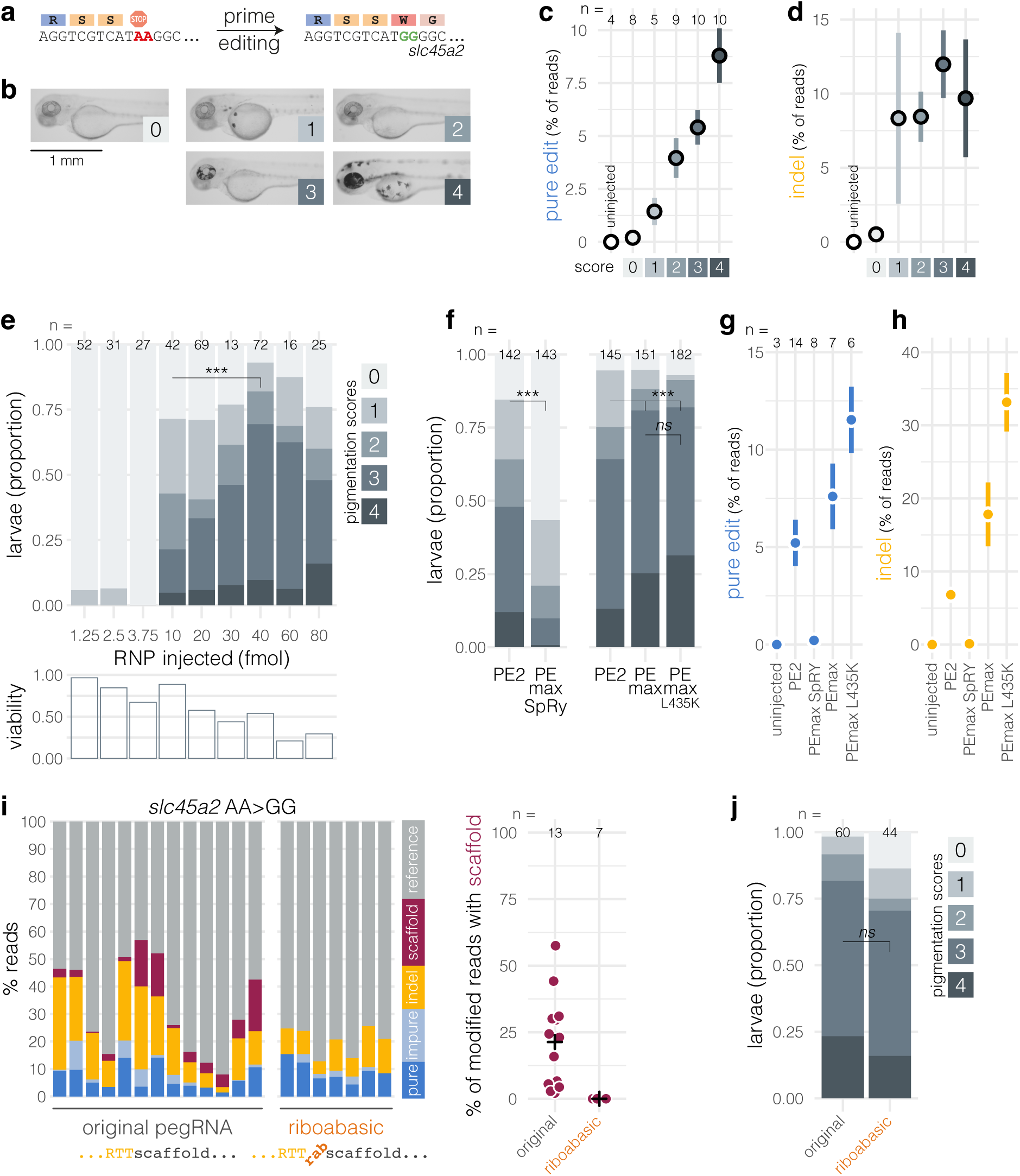
Optimising zebrafish prime editing using a pigmentation-based assay. a, Prime editing is used to create a two-nucleotide substitution in the pigmentation gene *slc45a2*, reverting a premature stop codon (TAA) into the wild-type tryptophan codon (TGG). **b,** Pictures of 3-dpf *slc45a2^W121*^* homozygous larvae injected with PE2/pegRNA RNP as examples of the pigmentation scoring method. **c,** Pure edit rates (% of reads, mean ± SEM) of larvae assigned to each pigmentation score. **d,** Indel rates (% of reads, mean ± SEM) of larvae assigned to each pigmentation score. **e,** (top) Phenotypic penetrance (proportion of larvae assigned to each pigmentation score) when gradually more PE2/pegRNA RNP was injected at the 1:1 ratio. Doses ≥10 fmol generally correlated with higher rates of pigmented larvae, in line with a previous report ^12^. 10 vs. 40 fmol: *** p < 0.001 by Fisher’s exact test. (bottom) Viability as proportion of larvae assessed as viable at 3 dpf, normalised to uninjected embryos. Doses >40 fmol lowered viability without further increasing rates of pigmented larvae. **f,** (top) Phenotypic penetrance (pigmentation scores) in larvae injected with 30 fmol of PE2, PEmax SpRY, PEmax ^16^, or PEmax^L435K^ RNP ^12^. ns p = 0.11, *** p < 0.001 by Fisher’s exact test. **g,** Pure edit rates (% of reads, mean ± SEM) of larvae injected with each PE RNP. **h,** Indel rates (% of reads, mean ± SEM) of larvae injected with each PE RNP. **i,** (left) Sequencing results from larvae injected with 30 fmol PEmax complexed with the original *slc45a2* AA>GG pegRNA or with the same pegRNA but carrying a riboabasic spacer (rab) between the RTT and the scaffold. Each bar represents an individual larva, with colours representing the percentage of reads assigned to each label. (right) Each larva’s scaffold incorporation rate as percentage of all modified reads (any read that is not reference). Black crosses mark the group means. Minimum sequencing depth was 364×. n = 20 samples. The original pegRNA group includes five PEmax samples also plotted in Supplementary Fig. 4. Only samples with ≥50 modified reads were included to calculate scaffold incorporation rates from a representative number of modified reads. **j**, Phenotypic penetrance (pigmentation scores) in larvae injected with PEmax RNP carrying the original *slc45a2* AA>GG pegRNA or the riboabasic spacer pegRNA, as in i. ns p = 0.11 by Fisher’s exact test.

Using our pigmentation recovery assay, we set out to calibrate injection conditions of the prime editor (PE) RNP in zebrafish. We optimised the ratio of PE protein to pegRNA (Supplementary Fig. 3a), the total dose of RNP injected (Fig. 1e), the site of injection (yolk or cell; Supplementary Fig. 3b), and the incubation temperature post-injection (Supplementary Fig. 3c). Maximal edit rates (∼5%) were obtained by injecting into the yolk ∼40 fmol of PE2 protein/pegRNA preassembled at the one-to-one ratio followed by incubation in a standard 28.5°C incubator.

Next, we tested PE protein variants: PEmax, PEmax^L435K^, and PEmax SpRY, which recognises any NRN (R = G or A) PAM rather than the standard NGG PAM ^12,16,17^. PEmax SpRY had much lower activity than PE2, reducing five-fold the proportion of highly pigmented larvae (score 3 and 4) (Fig. 1f). In contrast, PEmax and PEmax^L435K^ both increased 1.3-fold the proportion of highly pigmented larvae compared to PE2 (Fig. 1f). Deep sequencing broadly confirmed the phenotypic results (Fig. 1g and Supplementary Fig. 4): PE2 achieved 5 ± 4% pure edit rate; PEmax SpRY reduced this rate to 0.2 ± 0.3%; PEmax increased it to 8 ± 4% and PEmax^L435K^ to 12 ± 4%. Unfortunately, PEmax and PEmax^L435K^ concomitantly increased unwanted indels (Fig. 1h), from 7 ± 4% with PE2 to 18 ± 12% for PEmax and 33 ± 10% for PEmax^L435K^. To maximise editing rate, we conclude that PEmax or PEmax^L435K^ should be used.

The pegRNA’s 3′ end (PBS side) protrudes outside of the PE RNP, exposing it to exonuclease activity ^18^. We tested a chemically modified 3′-polyuridine tail optimised to protect the pegRNA end while also allowing the binding of La, an endogenous RNA-binding protein found to enhance prime editing ^19^. We also tested a longer pegRNA scaffold found to increase prime editing activity by 1.25-fold in HEK293T cells ^20,21^ (Supplementary Fig. 5a). Neither the alternative scaffold nor the optimised 3′-end pegRNA substantially modified the frequency of pigmented larvae (Supplementary Fig. 5b), suggesting that 3′-end degradation is not a major concern in zebrafish.

Insertion of a riboabasic spacer—a ribose sugar without a base—between the pegRNA RTT and scaffold prevented scaffold incorporations in a nuclease-based prime-editing strategy but has not been tested with standard prime editing in zebrafish ^13,22^. With the original *slc45a2* AA>GG pegRNA, 21 ± 17% of modified reads (any read that is not reference) carried scaffold-templated nucleotides (Fig. 1i). The riboabasic spacer completely eliminated scaffold incorporations (0 ± 0%) without substantially affecting editing efficiency (70% highly pigmented larvae, vs. 82% for the original pegRNA, Fig. 1j).

For a given edit and spacer sequence, varying PBS and RTT lengths gives hundreds of possible pegRNAs. To predict the optimal PBS length, it was suggested that the PBS:genome duplex Tm should approach 37°C ^14^. Reducing our *slc45a2* pegRNA PBS from 13 to 7 nt lowered its predicted Tm from 64°C to 37°C but decreased the rate of highly pigmented larvae from 33 to 7%, suggesting the Tm rule does not universally predict optimal PBS length (Supplementary Fig. 6a,b). As rational design is difficult, machine learning tools have been trained on human cell data to predict optimal pegRNA designs. To assess these tools in zebrafish, we compiled published edit rates from 39 pegRNAs encoding 31 edits across 19 loci ^5,6^ and tested 13 models from DeepPrime, PRIDICT, and OptiPrime ^21,23–25^. None produced a significant positive correlation between *in-vivo* edit rates and predicted pegRNA scores, indicating that these tools do not translate to zebrafish embryos (Supplementary Fig. 6c).

### Indels generated during prime editing in zebrafish show signature of microhomology-mediated end-joining

Prime editing relies on single-strand breaks and only generates low levels of unwanted indels in human cells ^5,11^. In zebrafish, however, prime editing generates surprisingly high indel rates ^5^. For example, PE2 generated 1.7-fold more unwanted indels than pure edits at *slc45a2* (Fig. 2a). 81% of these indels were deletions, the other 19% were insertions (Fig. 2b). To test whether the high indel rate was specific to *slc45a2*, we tested a G>A substitution (S33L) in *ctnnb1*: PEmax generated 37 ± 18% indels, 5.7-fold more than pure edits (6 ± 5%; Supplementary Fig. 7a). The majority of these indels were again deletions (Supplementary Fig. 7c). At both loci, indels were most frequent at the nick position, 3 bp upstream of the PAM. Indels were also frequent at the end of the RTT-templated nucleotides, which becomes a secondary nick position after RT synthesis and annealing of the 3′ flap generated by reverse transcription (Fig. 2c and Supplementary Fig. 7b). The high indel rates were previously reported so are not specific to PEmax or our injection conditions ^5^.

**Figure 2.**
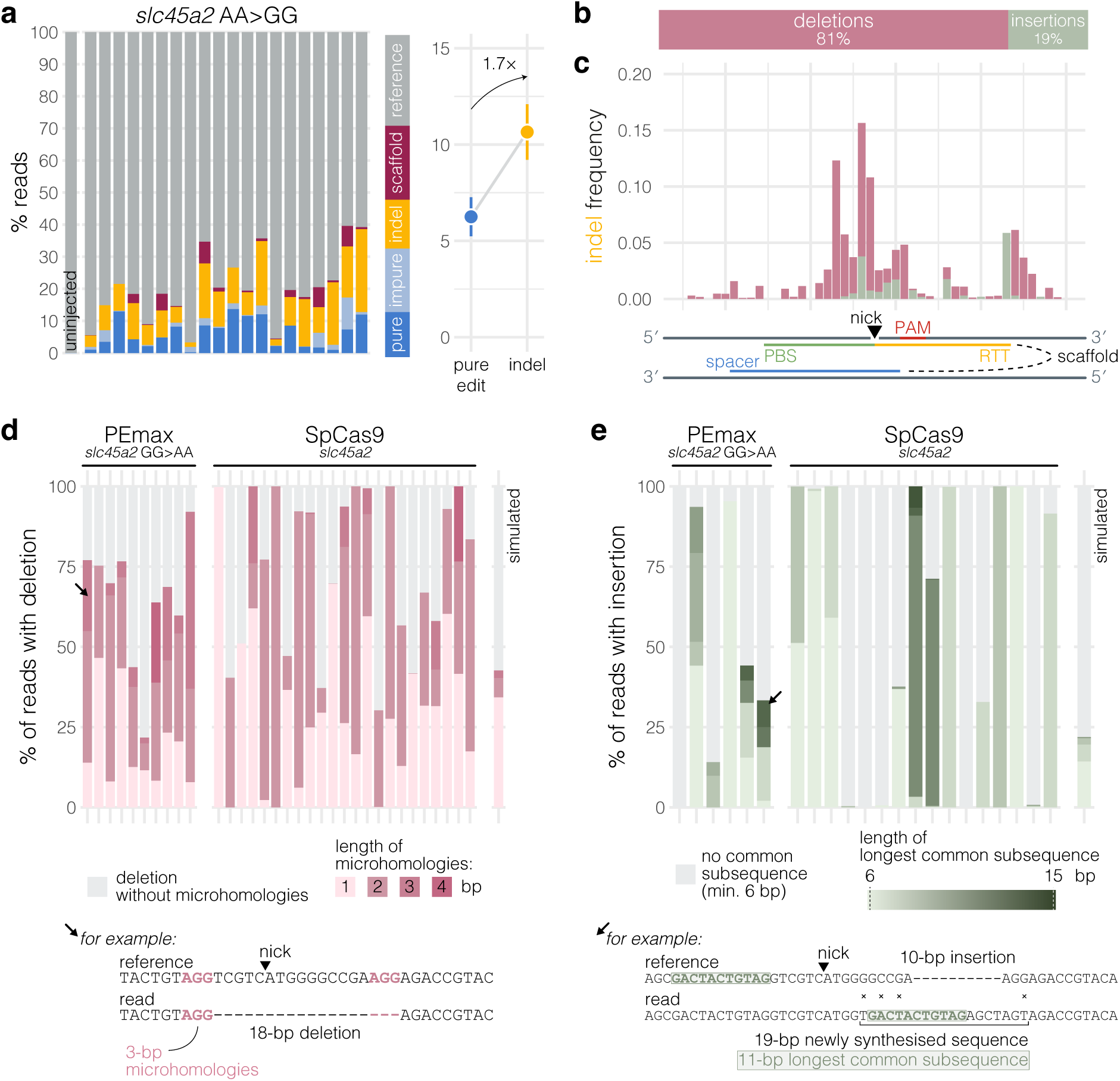
Indels generated during prime editing in zebrafish show signature of microhomology-mediated end-joining. a, (left) Sequencing results from larvae injected with PE2/pegRNA RNP. Each bar represents an individual larva, with colours representing the percentage of reads assigned to each label. Four samples with <10 modified reads were excluded. Minimum sequencing depth was 1377×. n = 20 injected samples. (right) The pure edit rate compared to the indel rate (mean ± SEM). **b,** In the average *slc45a2* AA>GG prime-edited sample, 81% of reads with an indel had a deletion, the other 19% had an insertion. **c,** Frequency of prime editing-generated indels across genomic positions at the *slc45a2^W121*^* locus. Samples included were larvae injected with the *slc45a2* AA>GG pegRNA complexed with PE2 or PEmax protein, for a total of n = 31 samples. Each frequency represents the average frequency across larvae of indels starting at this position. Each indel (n = 343 unique indels observed) contributes to the frequency once at its start position, which is the most 5′ nucleotide involved. For example, a 10-bp deletion which deleted nucleotide #90–99 is counted at #90. Scaffold incorporations are not included. **d,** (top) Of all sequencing reads with a deletion, percentage which showed microhomologies flanking the deletion. Each bar represents an individual larva injected with PEmax or SpCas9 RNP targeting the *slc45a2* locus. Darker pink colours represent longer microhomologies, grey represents deletions without microhomologies. Only samples with ≥30 reads with a deletion were analysed. There were 36–8030 reads with a deletion per sample included. The “simulated” sample represents 1000 simulated deletions. (bottom) Example of an 18-bp deletion flanked by 3-bp microhomologies; the arrow indicates the sample in which it was found. **e**, (top) Of all sequencing reads with an insertion, percentage which had a ≥6 bp continuous match between the “newly synthesised sequence” and the neighbouring sequences. Insertions generated during DSB repair in zebrafish are often accompanied with a multi-bp substitution (see example at bottom), they together form the “newly synthesised sequence” which is compared to the neighbouring sequences to find a possible match. Darker green colours represent longer matches between the newly synthesised sequence and the neighbouring sequences, grey represents newly synthesised sequences with a <6 bp match; that is, whose length was too short to analyse, or were untemplated, or used a template that was further away. Only samples with ≥30 reads with an insertion were analysed. There were 45–2866 reads with an insertion per sample included. The “simulated” sample represents 1000 newly synthesised sequences generated at random. (bottom) Example of a 19-bp newly synthesised sequence with at least 11 bp templated from a neighbouring sequence (green); the arrow indicates the sample in which it was found.

DSBs can be repaired by three main pathways: homology-directed repair (HDR), non-homologous end-joining (NHEJ), and microhomology-mediated end-joining (MMEJ). MMEJ is a very error-prone repair mechanism during which the helicase/DNA polymerase Polθ—encoded in zebrafish by *polq*—anneals short (≥1 bp) identical sequences on either side of the break (i.e., microhomologies) and fill-in the gaps, typically resulting in a deletion (Supplementary Fig. 8a,b). Early zebrafish embryos almost exclusively use MMEJ to repair DSBs, and seemingly cannot use HDR or NHEJ as back-up if MMEJ is unavailable. For example, *polq* knockout embryos do not survive Cas9 injections or doses of ionising radiation that are safe for wild-type embryos ^7,8^.

After prime editing at *slc45a2* in wild-type embryos (GG>AA, Supplementary Fig. 9a), we noticed that the majority of unwanted deletions were flanked by microhomologies (65 ± 20% vs. 43% of simulated deletions at the locus, Fig. 2d), consistent with repair of DSBs by MMEJ. Indeed, 75 ± 25% of deletions after SpCas9 DSBs at the same locus (same spacer sequence) were also flanked by microhomologies (Fig. 2d and Supplementary Fig. 9b), in line with zebrafish embryos relying heavily on MMEJ for DSB repair ^7,8^. Polθ can also create insertions haphazardly templated from sequences around the break ^26–28^. Half of the unwanted insertions generated during prime editing were templated from neighbouring sequences (56 ± 37% of insertion reads had a continuous match ≥6 bp between the inserted sequence and the neighbouring sequences vs. 22% of simulated insertions at the locus; Fig. 2e). The longest continuous match was 13 bp, while no match > 9 bp was observed in 1000 simulated insertions. This was again similar to insertions generated during repair of SpCas9 DSBs at the same locus (67 ± 42% of insertions were templated). We repeated this analysis on larvae prime-edited at *ctnnb1*: 81 ± 11% of unwanted deletions were flanked by microhomologies compared to 39% of simulated deletions (Supplementary Fig. 7d) and 50 ± 20% of insertions were templated compared to 22% of simulated insertions (Supplementary Fig. 7e). In summary, the majority of unwanted indels caused by prime editing in zebrafish show signatures of MMEJ: ∼75% of deletions are flanked by microhomologies and ∼50% of insertions are templated from neighbouring sequences.

To test whether this was specific to zebrafish, we reanalysed sequencing data from mouse embryos ^29^, focusing on two loci where PE2/PEmax caused indels (Supplementary Fig. 10a). Similar to zebrafish, 77 ± 36% of unwanted deletions were flanked by microhomologies, compared to ∼43% of simulated deletions (Supplementary Fig. 10b). We also found evident examples of templated insertions (Supplementary Fig. 10c). As MMEJ repairs DSBs by annealing microhomologies on opposite genomic strands (Supplementary Fig. 8a,b), we conclude that prime editing, even though it uses a Cas9 nickase, likely causes DSBs.

### Nicks are converted to DSBs endogenously

We then explored the potential source of these DSBs. In SpCas9, the HNH domain cuts the target strand (strand on which the gRNA is annealed) and the RuvC domain cuts the opposite non-target strand (Fig. 3b). The D10A mutation inactivates RuvC, so the resulting D10A nickase only cleaves the target strand. The H840A mutation inactivates HNH, so the resulting H840A nickase—used in prime editing—only cleaves the non-target strand. However, the H840A nickase reportedly retains residual nuclease activity ^30^. We first hypothesised that this was the source of the DSBs, which are then repaired by MMEJ. As indels were frequent, H840A nickase’s nuclease activity should be substantial, and therefore easily detectable. To test this *in vitro*, we incubated for 1 hr SpCas9, H840A nickase, or PEmax, with a double-stranded DNA substrate where each strand is fluorescently labelled to allow independent detection by denaturing PAGE (Fig. 3a). As expected, the non-target strand (strand opposite the gRNA) was cleaved by all three proteins. In contrast, only SpCas9 cleaved the target strand; cleavage by H840A nickase or PEmax was undetectable. We conclude that any residual nuclease activity of PEmax does not explain the high indel rates we observed during prime editing.

**Figure 3.**
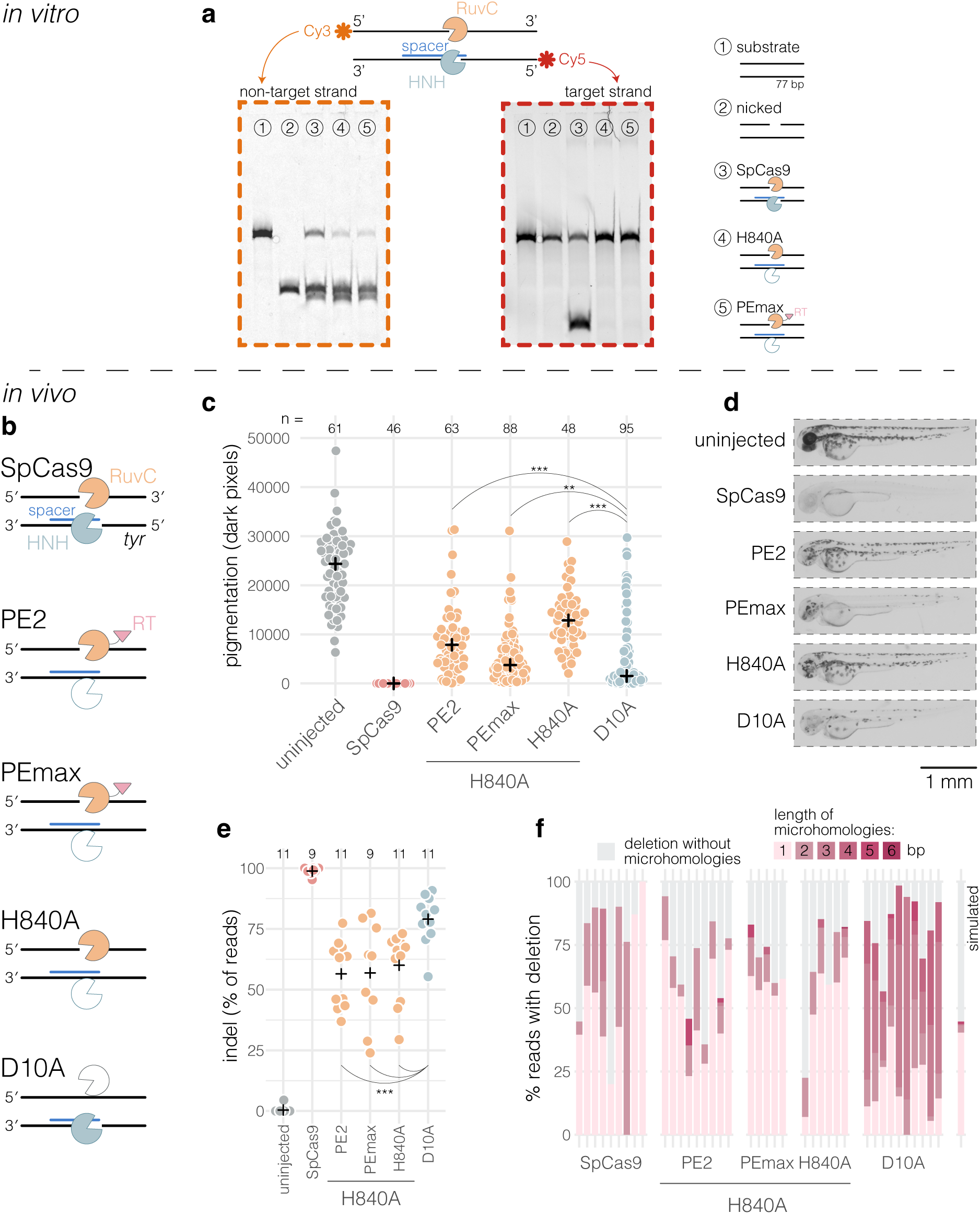
Nicks are converted to DSBs endogenously. a, *In-vitro* cleavage of a fluorescently-labelled double-stranded DNA substrate encoding a portion of *BFP* by SpCas9, H840A nickase, or PEmax, followed by denaturing PAGE. (left) Gel illuminated to detect Cy3 fluorescence emitted from the 5′–3′ forward strand. (right) Gel illuminated to detect Cy5 fluorescence emitted from the 3′–5′ reverse strand. **b,** At the *tyr* locus tested in zebrafish, SpCas9 cleaves both strands; the H840A nickases, which include PE2 and PEmax, cleave the 5′–3′ forward genomic strand; and the D10A nickase cleaves the 3′–5′ reverse genomic strand. **c,** Pigmentation of uninjected and injected larvae. Each dot represents the number of dark pixels per larva measured on pictures like in d. Black crosses mark the group medians. ** p = 0.005, *** p < 0.001 by Mann-Whitney U tests. **d,** Example pictures of 2-dpf uninjected and injected larvae. **e,** Indel rates (% of reads) of larvae injected with RNP made of each protein and a gRNA targeting *tyr*. Each dot represents an individual larva. Black crosses mark the group means. D10A vs. PE2 or PEmax or H840A: *** p < 0.001 by Welch’s t-tests. **f,** Of all sequencing reads with a deletion, percentage which showed microhomologies flanking the deletion. Each bar represents an individual larva. Darker pink colours represent longer microhomologies, grey represents deletions without microhomologies. Only samples with ≥50 reads with a deletion were analysed. There were 50–735 reads with a deletion per sample included. The “simulated” sample represents 1000 simulated deletions.

The D10A nickase reportedly generates much fewer DSBs than H840A nickase ^30^. If most prime-editing indels were indeed caused by residual PEmax nuclease activity, the D10A nickase should therefore not cause MMEJ indels in zebrafish embryos. To test this, we targeted the pigmentation gene *tyr* using SpCas9, PE2, PEmax, H840A nickase, or D10A nickase (Fig. 3b) ^31^. SpCas9-injected embryos were all unpigmented and 99 ± 1% of reads carried an indel (Fig. 3c–e). The three H840A nickases —PE2, PEmax, and H840A nickase—reduced pigmentation by 42–85% and led to mutations in about half of genome copies (56–60% indel rate). The D10A nickase was in fact more mutagenic than the H840A nickases, reducing pigmentation by 94% and pushing indel rate to 79 ± 10%. This is in stark contrast to D10A nickase activity in mammalian cells where it rarely causes indels ^26,32^. The HNH domain—the one still active in the D10A nickase—is known to cleave more than the RuvC domain ^33^, potentially explaining why D10A nickase was more mutagenic than H840A nickase in our experiment. Every protein caused a majority of microhomology-flanked deletions (Fig. 3f), from 66% of deletion reads for PE2 to 83% for D10A nickase. To test whether these observations also applied to base editing, which relies on target-strand cleavage by D10A nickase, we reanalysed a published experiment that tested the base editor AncBE4max ^9,34^. AncBE4max generated 22 ± 5% unwanted indels (Supplementary Fig. 11a). In line with our results, 75 ± 4% of the deletions were flanked by microhomologies compared to 48% of simulated deletions (Supplementary Fig. 11b).

As target-strand cleavage by PEmax was undetectable *in vitro* and that D10A nickase also caused microhomology-flanked deletions in zebrafish embryos, we conclude that most MMEJ-repaired DSBs were not directly made by PEmax. Moreover, H840A nickase without RT and PEmax/gRNA both generated MMEJ indels, excluding an indel-causing mechanism involving the RT or the prime-editing flaps ^35^. Instead, we propose that nicks are converted into DSBs endogenously by replication forks. When a replication fork arrives at a nick, daughter strand synthesis is interrupted because the template strand is broken, leading to a single-ended DSB (seDSB) (Supplementary Fig. 13) ^36–40^. Canonical MMEJ operates on double-ended DSBs, but an alternative pathway termed fork-MMEJ was recently described to repair replication-associated seDSBs, producing longer, asymmetric deletions ^41^. Across four loci, Cas9 nickase activity indeed produced longer deletions than repair of SpCas9 DSBs (deletions >10 bp were 2–9× more frequent after Cas9 nickase vs. after SpCas9; Supplementary Fig. 12a). At *tyr*, while SpCas9 DSBs produced short deletions that were mostly symmetric, deletions found after Cas9 nickase activity were obviously asymmetric, extending 3′ of the cut when the forward 5′–3′ genomic strand was cleaved, and 5′ of the cut when the reverse 3′–5′ genomic strand was cleaved (Supplementary Fig. 12b,c). These observations suggest that zebrafish embryos frequently use fork-MMEJ to repair replication-associated seDSBs (Supplementary Fig. 13).

### Prime editing in the absence of maternally deposited Polθ achieves near-perfect precision

As most prime-editing indels seemed created by MMEJ, we wondered whether inhibition of MMEJ would reduce indels. Polθ, encoded in zebrafish by *polq*, is essential to MMEJ but homozygous knockout (KO) zebrafish are fertile ^7,8^. As in virtually all metazoans, early zebrafish embryonic development up to the maternal-to-zygotic transition relies on mRNA and proteins that were maternally deposited in the oocyte ^42^. As mRNA of *polq* and other MMEJ factors are maternally deposited (Supplementary Fig. 14) ^7^, we used maternal *polq* mutant embryos to prevent MMEJ early in development. We tested a one-bp substitution (G>C in *cacng2b*) in wild-type embryos, maternal-only heterozygous mutants (offspring of an outcross homozygous *polq* KO mother × wild-type father), and maternal-zygotic homozygous *polq* mutants (offspring of a homozygous *polq* KO incross) (Fig. 4a). In wild-type embryos, pure edit rate was 13 ± 6% but indel rate was 1.6-fold higher (21 ± 11%). Strikingly, indels were completely absent in maternal-zygotic *polq* mutants (0 ± 0%) and rare in maternal-only mutants (0.7 ± 1.2%). The pure edit rate also surged in *polq* mutants, reaching 45 ± 25% in maternal-only mutants and 53 ± 28% in maternal-zygotic mutants.

**Figure 4.**
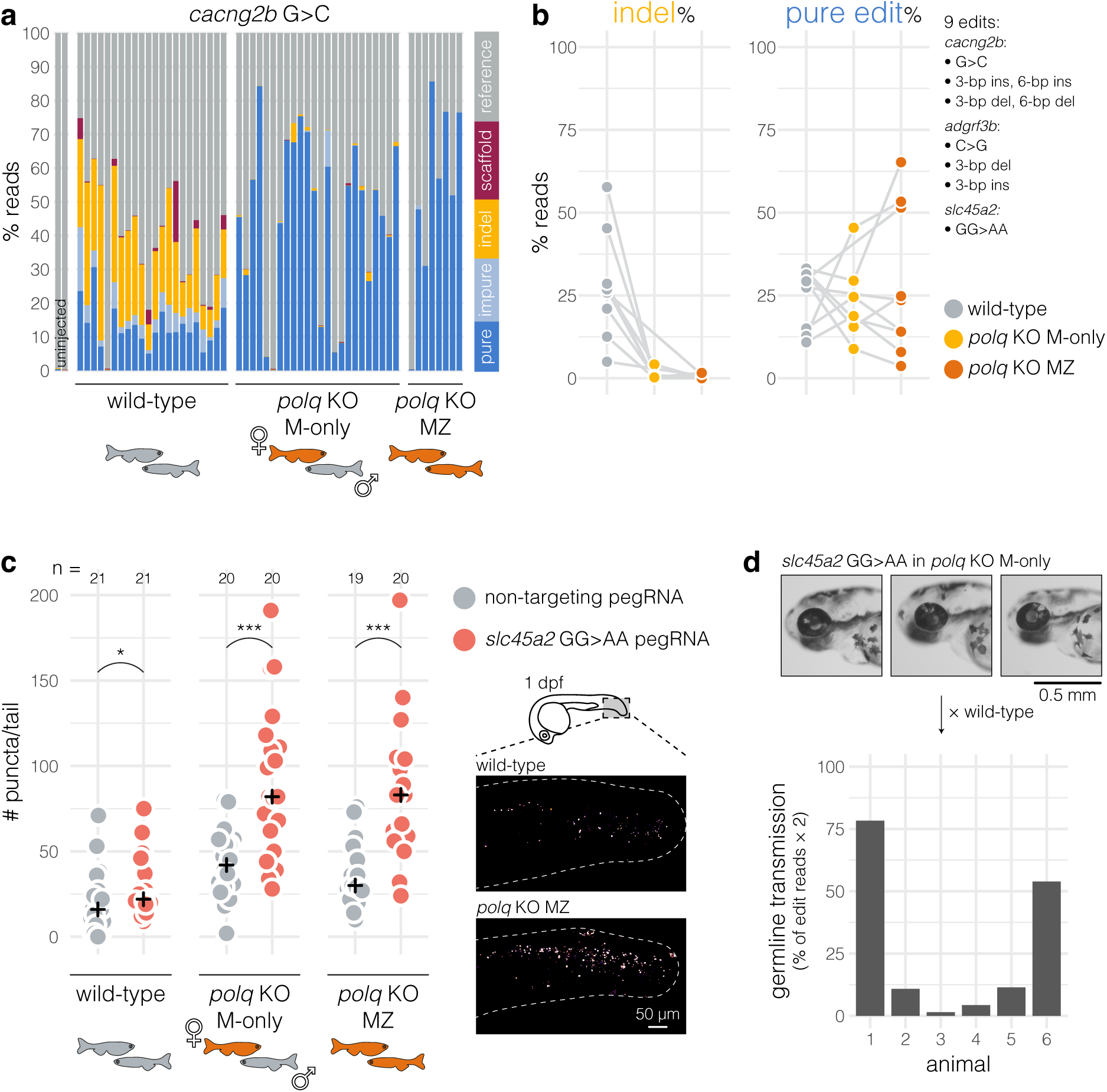
Prime editing in the absence of maternally deposited Polθ achieves near-perfect precision. a, Sequencing results from wild-type, *polq* maternal-only heterozygous KO, and *polq* maternal-zygotic homozygous KO larvae uninjected or injected with PEmax complexed with a pegRNA encoding a one-bp substitution (G>C) in *cacng2b*. Each bar represents an individual larva, with colours representing the percentage of reads assigned to each label. Cartoons below represent the parental crosses that generated the injected embryos; orange fish are homozygous *polq* KO. Minimum sequencing depth was 60×. n = 54 injected samples. KO, knockout; M-only, maternal-only; MZ, maternal-zygotic. **b,** Each datapoint represents the average indel (left) or pure edit (right) rate (% of reads) of 7–24 larvae of same genotype (wild-type, *polq* maternal-only heterozygous KO, or *polq* maternal-zygotic homozygous KO) and injected with the same pegRNA. Lines connect datapoints from the same pegRNA. n = 9 pegRNAs (7 for maternal-only *polq* KO; *adgrf3b* 3-bp insertion and *cacng2b* 6-bp deletion are missing); n = 278 injected samples in total. **c,** Detection of apoptotic cells using acridine orange staining. (left) Each datapoint represents the number of acridine-positive puncta in the tail of an individual 1-dpf wild-type, *polq* maternal-only heterozygous KO, or *polq* maternal-zygotic homozygous KO embryo injected with PEmax complexed with a non-targeting pegRNA or the *slc45a2* GG>AA pegRNA. Black crosses mark the group medians. Cartoons below represent the parental crosses that generated the injected embryos; orange fish are homozygous *polq* KO. * p = 0.011, *** p < 0.001 by Mann-Whitney U tests. (right) Example images of the tail of a 1-dpf wild-type or *polq* maternal-zygotic homozygous KO embryo injected with the *slc45a2* GG>AA pegRNA showing acridine orange-positive puncta. **d,** (top) Example pictures of 3-dpf *polq* maternal-only heterozygous KO larvae injected with PEmax^L435K^/*slc45a2* GG>AA pegRNA that were selected to be grown because they had patches of eye depigmentation. (bottom) Germline transmission rates of the *slc45a2* GG>AA edit measured from six F1 clutches. Minimum sequencing depth was 2283×.

We tested a total of 9 edits across three loci (Supplementary Fig. 15): three substitutions (1 or 2 bp, including *cacng2b* G>C), three short insertions (3 or 6 bp), and three short deletions (3 or 6 bp). Total absence of Polθ abolished nearly all indels (Fig. 4b; maximum 1.6 ± 1.0% for *slc45a2* GG>AA in maternal-zygotic KO). The dependence on Polθ to generate deletions was indeed close to absolute: we only found 11 deletion alleles in 31,568 sequencing reads; 8 of which had no or 1-bp microhomologies, which may be expected by chance (Supplementary Fig. 16). Similarly, there were just 8 insertion alleles left, all 1–6 bp, demonstrating that all templated insertions were dependent on Polθ, not on the RT as previously hypothesised ^5^. Maternal-only mutants had similarly low indel rates (maximum 4 ± 3% for *adgrf3b* C>G, all others <1%). Interestingly, absence of Polθ also eliminated scaffold incorporations at *slc45a2* (4 ± 5% in wild types, 0 ± 0% in n = 21 *polq* mutants, see Supplementary Note 1). Effects on edit rates were often drastic too (Fig. 4b): for 3/9 edits, pure edit rates raised >50% in maternal-zygotic KO, with almost half of samples (11/28) even reaching >70% pure edits. Pure edit rates were largely unchanged for 4/9 edits (<15% difference) but declined for 2/9 (−18, −26%). Maternal-only mutants gave similar edit rates as maternal-zygotic ones (6/7 edits with <15% difference with maternal-zygotic KO). Using heterozygous (maternal-only) *polq* mutants thus provides most benefits from MMEJ inhibition while allowing edited animals in wild-type background to be created in the F1.

Without Polθ, are DSBs repaired by a mechanism that is far less mutagenic than MMEJ (e.g., NHEJ or HDR) or are DSBs simply left unrepaired? If the latter is true, cells harbouring a DSB should eventually enter apoptosis. We used acridine orange to stain apoptotic cells in wild-type, maternal-zygotic, and maternal-only *polq* mutant embryos injected with a non-targeting pegRNA or with the *slc45a2* GG>AA pegRNA (Fig. 4c). Prime editing in wild-type embryos, independently of injections, caused some apoptosis as the number of acridine-positive puncta increased slightly (1.4-fold) compared to the non-targeting pegRNA-injected embryos. This effect was stronger in absence of maternal Polθ, with apoptotic cells rising 2-fold in maternal-only mutants and 2.8-fold in maternal-zygotic mutants. To compare these to other forms of DNA damage, we exposed wild-type embryos to cadmium, a mutagenic heavy metal; AOH1160, a PCNA inhibitor; or UV radiation. The number of acridine-positive puncta increased from 4 to 12-fold (Supplementary Fig. 18). We propose that prime-editing nicks are converted to DSBs which are left unrepaired when maternal Polθ is not available, pushing cells harbouring a DSB to enter apoptosis, eventually leaving only wild-type and correctly edited cells.

Finally, to confirm germline transmission of the prime edits in *polq* mutants, we injected maternal-only mutants with a PEmax^L435K^/*slc45a2* GG>AA RNP, grew larvae with patches of depigmentation, and sequenced their offspring (Fig. 4d). All six animals tested transmitted the GG>AA edit at rates 1.5–78%, confirming that prime editing in *polq* mutants can be used to establish stable lines.

In summary, almost every unwanted indel generated during prime editing in zebrafish embryos was created by Polθ-dependent MMEJ. Prime editing in *polq* knockouts achieves virtually perfect precision and can lead to edit rates >50%.

### Supplementary Note 1

Absence of Polθ eliminated scaffold incorporations at the *slc45a2* locus (4 ± 5% in wild-type larvae, 0 ± 0% in n = 21 *polq* mutants, Supplementary Fig. 15), but paradoxically our results indicate that the RT frequently proceeds past the end of the RTT without the riboabasic spacer, at least at the *slc45a2* locus (Fig. 1i). It may be that the RT regularly writes a scaffold-templated sequence in the edited 3′ flap, which is then installed in the genome as an insertion by Polθ taking the flap as template. In absence of Polθ, the edited 3′ flap may correctly anneal to the genome, pushing its scaffold-templated end as a flap that is then excised like the unedited 5′ flap. Some scaffold incorporations indeed looked like Polθ-mediated insertions (Supplementary Fig. 17), consisting of a patchwork of scaffold-templated sequences sometimes followed by sequences that exist earlier in the RTT. At the *slc45a2* locus, the end of the RTT-templated nucleotides matches the start of the scaffold (edited 3′ flap should end with GGAGA, first scaffold-templated nucleotides are expected to be GGA), which may facilitate the annealing of microhomologies. It remains to be seen whether absence of Polθ also eliminates scaffold incorporations at other loci.

## Discussion

The promise of nick-based genome editing like prime editing is to limit unwanted indels by avoiding DSBs. However, prime editing in zebrafish embryos is highly error-prone, in some cases generating as many indels as genome editing using Cas9 nuclease. We show that almost every prime-editing indel was created by Polθ-dependent MMEJ. Prime editing in *polq* knockouts achieves virtually perfect precision and can lead to edit rates >50%.

In human cell cultures, prime editing generates far fewer indels than in zebrafish embryos ^5,11^. In zebrafish, we show that these indels are created by MMEJ: most are microhomology-flanked deletions and templated insertions dependent on Polθ activity. MMEJ being a repair mechanism that requires two free DNA ends for microhomology search and annealing ^43^, it follows that prime editing causes DSBs in the zebrafish genome. We initially considered that the H840A nickase used in prime editing might be cleaving both strands ^30^, but this appeared unlikely as DSBs were undetectable *in vitro* and that the D10A nickase also caused MMEJ indels in zebrafish. The likely mechanism, first demonstrated in bacteria ^37^ and later in many species ^36,38–40^, involves the conversion of nicks into DSBs by replication forks. This model can explain the discrepancy with human cells as early zebrafish embryos have a cell cycle that is almost a hundred times faster than most mammalian cells (15 min vs. 24 hr). During the first few hours after fertilisation, every locus in the zebrafish genome must necessarily be visited by a replication fork every 15 min, creating a high chance of collision between nicks and replication forks. This principle probably applies across species and cell types: the likelihood of generating a DSB during nick-based genome editing may depend partly on cell cycle length.

When a replication fork collides with a nick, the resulting DSB is initially single-ended (seDSB; i.e., only one side is dsDNA). This seDSB can likely be repaired by fork-MMEJ ^41^, generating the long, asymmetric deletions we observed. The seDSB can also become a double-ended DSB (deDSB; i.e., dsDNA on both sides), the canonical substrate of MMEJ, through at least two mechanisms: another replication fork running in the opposite direction collides with the seDSB, or—if the nick was in the lagging strand—synthesis ‘bypassing’ the break by placing a new Okazaki fragment after ^36,40,44^. In an *in-vitro* experiment ^45^, POLθ enabled skipping of an ssDNA gap without requiring a DSB via a mechanism termed microhomology-mediated gap skipping (MMGS). In theory, MMGS creates deletions flanked by microhomologies, but it is unclear why these deletions would be asymmetric and longer than after repair of SpCas9 DSBs, while fork-MMEJ readily explains this. Fork-MMEJ is predicted to generate different mutations depending on direction of replication and whether the leading or lagging strand is cleaved, with some configurations even preventing repair. Exploring the role of fork-MMEJ in zebrafish embryos will therefore require mapping strong origins of replication, so that nicks can be made at loci where the direction of the replication fork is known.

Absence of maternally deposited *polq* mRNA/Polθ protein drastically reduced indel rates, even when the embryo inherited a functional *polq* gene from its father. This wild-type paternal copy becomes active after the maternal-to-zygotic transition around 5 hours post-fertilisation (hpf), but seemingly does not create many indels. This observation leaves two possible options: either every break is repaired within the first few hours and PEmax does not make new breaks after that; or repair of breaks does not leave indels after 5 hpf. The first option is unlikely as ample wild-type genome copies remain and that PEmax is probably still present at sufficient concentrations in cells to make breaks for many hours or even days after injection ^46^. For example, to test a photoactivable CRISPR system, Hu et al. (2025) injected one-cell stage embryos with 6 fmol of inactivated Cas9/gRNA RNP then exposed the larvae two days later with UV to activate the RNPs, which still caused indels in ∼35% of genome copies. After the 10^th^ cell division, which coincides with the start of the maternal-to-zygotic transition ^47^, cell cycle length slows down ^48,49^. This may explain why, after 5 hpf, new breaks by PEmax are repaired without leaving indels: replication forks are now less frequent, leaving enough time for the nick to be ligated before it is converted into a DSB. Accordingly, we predict that applications using prime editing later in development, for example for tissue-specific genome editing, would generate lower indel frequencies.

Finally, it is surprising that nicks in the zebrafish genome are so mutagenic. If the thousands of nicks naturally occurring in the first few hours of development were each causing an indel ^50^, about a hundred protein-coding genes would be inactivated at each generation, which is inconceivable. The present situation may be somewhat specific to Cas9 nickase. By remaining bound to the DNA ends after cleavage ^51^, Cas9 may block repair for long enough that a replication fork is likely to encounter the nick and convert it into a DSB before it is repaired accurately by a nick-specific pathway like single-strand break repair. Another nicking enzyme which does not ‘cling’ to DNA after cleavage, such as I-AniI homing endonuclease, could be tested to confirm this^52,53^.

As prime editing relies on nicks rather than DSBs, it is widely expected to cause minimal DSBs and unwanted indels. Our work presents a striking counterexample, the zebrafish embryo, where any Cas9 nickase activity causes high levels of indels. By identifying Polθ-dependent MMEJ as the sole culprit behind the unwanted prime-editing indels, our work also provides a simple solution: prime editing in MMEJ-deficient embryos is virtually error-free and can achieve edit rates >50%. We anticipate that this approach will further expand the utility of zebrafish as a model for genetic diseases and functional genomics. In the future, error-free prime editing may also prove useful for lineage tracing ^54^ and F0 phenotyping if edit rates can be raised further ^29^. Our finding that the D10A nickase also causes MMEJ indels suggests that precision of other nick-based genome editing methods, such as base editing and nickase-based HDR ^52^, could also benefit from Polθ inactivation. This may be true for many model organisms as prime editing also causes indels with signature of MMEJ in mouse embryos. *Drosophila*, *Xenopus*, and *C. elegans* embryos also have rapid cell cycles (respectively 9, 30, 15–50 min) ^55–57^ and can repair DSBs by MMEJ ^58–60^.

## Methods

### Animals

Adult zebrafish (*Danio rerio*) were reared in Institut de la Vision (Paris, France) fish facility at 25°C on a 14 hr:10 hr light:dark cycle with gradual lights-on and lights-off transitions to simulate sunrise and sunset. To obtain eggs, male and female zebrafish were isolated in breeding boxes overnight, separated by a divider. Around 10 AM the next day, the dividers were removed and eggs were collected. The embryos were then raised in 10-cm Petri dishes filled with egg medium (0.3 g/L Instant Ocean and 0.00005% methylene blue) in a 28.5°C incubator on a 14 hr:10 hr light:dark cycle. Debris and dead or dysmorphic embryos were removed every other day with a Pasteur pipette under a bright-field microscope and the fish water replaced. At the end of the experiments, larvae were euthanised with an overdose of tricaine (MS-222, Sigma-Aldrich).

All animal procedures were performed in accordance with French and European Union animal welfare guidelines and were approved by the Animal Experimentation Ethics Committee of Sorbonne Université (APAFIS#21323- 2019062416186982 and APAFIS #49266-2024050214115020) (number 005).

Wild types refer to TL (Tüpfel long fin) zebrafish. The *polq* KO line was generated by Carrara et al., 2025^7^. Using Sanger sequencing, we independently verified the mutation to be a 219-bp deletion spanning exon 6 and exon 7/32 (chr21:22,866,462–22,866,680 on reference genome GRCz11). We genotyped animals by running *polq* amplicons on a 1% agarose gel stained with ethidium bromide (Dutscher). PCR primer sequences were from Carrara et al., 2025 and are provided in Supplementary File 1. Maternal-zygotic homozygous *polq* KO refers to offspring of an incross where both parents were homozygous *polq* KO. Maternal-only heterozygous *polq* KO refers to offspring of an outcross where the mother was homozygous *polq* KO and the father was TL wild-type.

### Synthesis of PE proteins

We adapted a protocol for synthesis of SpCas9 ^61^. The protocols will be described in a subsequent publication (Alvarez Vargas et al., 2026. *in preparation*). Except for PE2, all PE proteins used in this work were based on the PEmax architecture ^16^. The SpRY variant included the mutations described by Walton et al., 2020 ^17^ except A61R. pET28-PE-His6 plasmid was introduced in Rosetta 2(DE3) competent cells (Sigma-Aldrich) for protein expression. A three-step purification was performed using HisTrap affinity chromatography followed by cation and anion exchange columns. PE protein stocks were prepared in 20 mM HEPES pH 7.5, 350 mM KCl, 1 mM TCEP, 10% glycerol at 60 µM for PE2, 60–90 µM for PEmax, 70 µM for PEmax SpRY, 90 µM for PEmax^L435K^. Stocks were stored at −80°C or−20°C before use.

### Preparation and injection of PE/pegRNA RNP

Synthetic pegRNAs were bought from Integrated DNA Technologies (IDT) as “Custom CRISPR gRNA” with standard desalting purification and without Alt-R modifications. Sequences are provided in Supplementary File 1. Each pegRNA was received as a pellet which was resuspended in Duplex buffer (IDT) to form a 100 µM stock. Aliquots of this stock were stored at −80°C before use. On the morning of injections, the pegRNA was mixed with the PE protein in equimolar proportions (e.g., 0.6 µL Duplex buffer; 0.9 µL pegRNA 100 µM; 1.5 µL PE protein 60 µM) and the mix was incubated at 37°C for 5 min then cooled on ice to generate a 30 µM PE/pegRNA RNP solution. Approximately 1 nL of this RNP solution (i.e., 30 fmol RNP) was injected into the yolk at the one-cell stage before cell inflation.

*slc45a2* AA>GG pegRNA with riboabasic spacer (“rSpacer”) used in Fig. 1i was bought from Horizon Discovery with HPLC purification. The PEmax/riboabasic pegRNA RNP was prepared and injected as above.

The spacer sequences of the *cacng2b* and *adgrf3b* pegRNAs were taken from Petri et al., 2025 ^5^. The *cacng2b* G>C pegRNA had the same sequence as the one used by Petri et al., 2025; others were modified. The following edits inactivated the PAM: *cacng2b* G>C, *cacng2b* 6-bp deletion, *adgrf3b* C>G, *adgrf3b* 3-bp deletion. The following edits did not inactivate the PAM: *slc45a2* AA>GG, *slc45a2* GG>AA, *ctnnb1* G>A, *cacng2b* 3-bp deletion, *cacng2b* 3-bp insertion, *cacng2b* 6-bp insertion, *adgrf3b* 3-bp insertion. All *cacng2b* and *adgrf3b* pegRNAs carried a 3′ polyuridine (UUUU) tract; others did not, except for the *slc45a2* pegRNA with optimised 3′ end (Supplementary Fig. 5a). pegRNA sequences can be found in Supplementary File 1.

In the experiment testing an excess of pegRNA (Supplementary Fig. 3a), the pegRNA was resuspended in Duplex buffer to form a 200 µM stock so that the pegRNA could be added in higher amounts.

In the experiment testing RNP doses (Fig. 1e), we prepared a 20 µM RNP solution as above and diluted some to make a 2.5 µM RNP solution by adding a 1:1 mix of Duplex buffer and protein buffer (see *Synthesis of PE proteins*). We then injected 0.5, 1.0, 1.5 nL of the 2.5 µM solution, which amounts to 1.25, 2.5, 3.75 fmol RNP; and 0.5, 1.0, 1.5, 2.0 nL of the 20 µM solution, which amounts to 10, 20, 30, 40 fmol RNP. In a separate experiment, we prepared a 40 µM RNP solution as above and injected 0.5, 1.0, 1.5, 2.0 nL, which amounts to 20, 40, 60, 80 fmol RNP. We also injected 3 nL (120 fmol RNP) of this solution but too few larvae (6/43) survived until 3 dpf to be analysed. For the common doses, results from both experiments replicated even though volume injected varied (less than ± 10% difference in proportion of highly pigmented larvae).

In the experiment testing different sites of injections (Supplementary Fig. 3b), eggs taken from the same batch were injected in the yolk; or in the single cell before inflation; or in the single cell after it inflated; or in both cells after the first division (1 nL in each cell).

### Pigmentation recovery assay

The *slc45a2^W121*^* mutant line was generated by base editing in our previous work ^9^. *slc45a2^W121*^* homozygous embryos were injected at the one-cell stage with PE RNP encoding the TAA(*)>TGG(W) edit to restore the wild-type (TGG) allele. At 3 dpf, each larva was scored on a scale from 0 to 5 mainly based on its eye pigmentation: score 0 if the larva was as unpigmented as *slc45a2^W121*^* homozygous larvae; score 1 if the eye was unpigmented but the larva had some pigmented cells elsewhere; score 2 if the eye had a single patch of pigmented cells; score 3 if more than approximately a third of the surface of the eye was pigmented; score 4 if most of the eye surface was pigmented. To avoid any bias in case one eye was pigmented but the other was not, only the right or left eye was scored during a given experiment. All scoring was done blinded to the condition. Dysmorphic larvae were scored if their eyes were sufficiently developed. Viability reports the number of larvae looking healthy at the end of the experiment (3–6 dpf) divided by the total number of injected embryos at 0 dpf; that is, including any unfertilised eggs or embryos damaged by the injection needle. Larvae that had not hatched naturally from the chorion at 3 dpf were counted as unviable. To exclude loss of viability unrelated to the experiment (e.g., unfertilised eggs), we divided this proportion to the viable proportion of uninjected embryos of the same experiment.

### Targeting *slc45a2* or *tyr* with SpCas9 or Cas9 nickases

The synthetic *tyr* gRNA was made of two components which were bought separately from IDT: the crRNA (Alt-R CRISPR-Cas9 crRNA) and tracrRNA (Alt-R CRISPR-Cas9 tracrRNA). The crRNA sequence was taken from our previous work ^1^. The crRNA and tracrRNA were received as pellets, which were individually resuspended in Duplex buffer (IDT, received with the tracrRNA) to form 200 µM stocks. Stocks of crRNA and tracrRNA were stored at −80 °C before use. The crRNA was annealed with the tracrRNA by mixing 3.7 µL crRNA 200 µM; 3.7 µL tracrRNA 200 µM; 4.6 µL Duplex buffer. The mix was heated to 95°C for 5 min, then cooled on ice, to obtain a 61 µM gRNA (crRNA/tracrRNA duplex) solution. For each protein (SpCas9, PE2, PEmax, H840A nickase, D10A nickase), 2 µL of the gRNA solution were mixed with 2 µL of protein at 60 or 61 µM. SpCas9 (Alt-R S.p. Cas9 Nuclease V3), H840A nickase (Alt-R S.p. Cas9 H840A Nickase V3), and D10A nickase (Alt-R S.p. Cas9 D10A Nickase V3) were bought from IDT. PE2 and PEmax proteins were synthesised in house (see *Synthesis of PE proteins*). Each mix was incubated at 37°C for 5 min then cooled on ice, generating six 30 µM RNP solutions. Approximately 1 nL of each solution (∼30 fmol RNP) was injected in the yolk at the one-cell stage. Targeting *slc45a2* with SpCas9 followed the same steps as above. The crRNA sequences are provided in Supplementary File 1.

For the experiment targeting *tyr*, we photographed the injected larvae at 2 dpf under a brightfield microscope (see *Pictures*). To speed up the process, we photographed multiple larvae in the same field of view. The images were then segmented into pictures of individual larvae using a YOLO v8 (ultralytics v8.3.7) model ^62^ trained on 350 images manually annotated using <u>app.roboflow.com</u>. We then counted, for each picture, the number of pixels with grey value ≤116 (0 is black, 255 is white) using the R package ijtiff, returning the number of dark pixels plotted in Fig. 3c.

### Germline transmission of *slc45a2* GG>AA

Maternal-only heterozygous *polq* KO embryos were injected at the one-cell stage with 30 fmol of PEmax^L453K^/*slc45a2* GG>AA pegRNA RNP. At 3 dpf, larvae with patches of eye depigmentation visible under a brightfield microscope were selected to be grown. Adults were then crossed to wild-type animals and we extracted genomic DNA from pools of 20–40 F1 embryos (see *Preparation of samples for deep sequencing*). From the genomic DNA, we amplified a 546-bp amplicon spanning exon 1/7 of *slc45a2* (primer sequences in Supplementary File 1). Amplicons were sent to Plasmidsaurus for Nanopore sequencing (PCR Premium Standard service). We aligned the fastq reads to the reference amplicon sequence using minimap2 v2.22. To calculate the rate of GG>AA edit, we counted the number of AA reads in IGV and divided this count by the total number of reads at this position. We also calculated edit rates using the CRISPRLungo web interface ^63^ and found that both methods gave a <1% difference in edit rates for every sample. As the F0 injected animals were crossed to wild types to generate the F1 offspring sequenced, the true germline transmission rate is expected to be double the edit rates measured in the F1; Fig. 4d therefore reports double the measured edit rates.

### Preparation of samples for deep sequencing

Individual larvae were anaesthetised with tricaine (MS-222) and their genomic DNA extracted by HotSHOT ^64^ as follows. Individual larvae were transferred to a 96-well PCR plate or strips of 8 tubes. Excess liquid was removed from each well before adding 50 µL of 1× base solution (25 mM KOH, 0.2 mM EDTA in water). Plates were sealed and incubated at 95°C for 30 min then cooled to room temperature before the addition of 50 µL of 1× neutralisation solution (40 mM Tris-HCl in water). Genomic DNA was then stored at –20°C.

PCR primers were designed for each target locus using Primer-BLAST (NCBI) to amplify a window of 150–200 bp with at least 20 bp between each primer binding site and the predicted cut site. PCR primers were ordered with a Nextera overhang at the 5′ end of each primer to allow indexing (see Supplementary File 1). Some PCR primer sequences were taken from previous studies ^1,5^.

Each PCR well contained: 5.13 µL dH2O, 1.8 µL 5× Phusion HF buffer (New England Biolabs), 0.3 µL DMSO, 0.18 µL dNTP (10 mM), 0.45 µL forward primer (10 µM), 0.45 µL reverse primer (10 µM), 0.09 µL Phusion High-Fidelity DNA Polymerase (New England Biolabs), 1.0 µL genomic DNA; for a total of 9.4 µL. The PCR plate was sealed and placed in a thermocycler. The PCR program was: 98°C – 5 min, then 40 cycles of: 98°C – 10 sec, 60°C – 30 sec, 72°C – 30 sec, then 72°C – 10 min, then cooled to 10°C until collection. The PCR product’s concentration was quantified with Qubit (dsDNA Broad Range Assay) and its length was verified on a 1% agarose gel with ethidium bromide (Dutscher). Excess primers and dNTPs were removed by ExoSAP-IT (Thermo Fisher) following the manufacturer’s instructions. The samples were then sent for Illumina MiSeq, which used MiSeq Reagent Nano Kit v2 (300 Cycles).

### Sequencing analysis of prime-editing samples

Illumina MiSeq data was received as two fastq files for each well, one forward and one reverse. The paired-end reads were aligned to the reference amplicon with bwa v0.7.18 and the resulting bam alignment file was sorted and indexed with samtools v1.6 ^65^. Alignments were then filtered to keep only reads with less than 20% of their length soft-clipped and spanning at least 90 bp of the reference sequence and 30 bp on each side of the predicted cut site. To check the alignment and filtering steps, bam alignment files were visualised with IGV v2.16.2. The resulting filtered bam file were converted back to forward and reverse fastq files using bedtools v2.30.0 ^66^. These filtered fastq files were used as input to CRISPResso2 v2.2.15 ^67^ ran in command line. CRISPResso2 was ran for each pair of fastq files in standard amplicon mode—not in prime-editing mode—to produce alignments. From CRISPResso2’s output, each *Alleles_frequency_table.txt* file was used as input for our custom “CUTTER” (the CRISPR mutations caller) R package available at <u>github.com/francoiskroll/cutter</u>. Briefly, function callMutations(…) produced a csv table listing all mutations on individual reads by calling insertions, deletions, and substitutions for each alignment. Any mutation that was found in fewer than three reads or was not within ± 12 bp of the cut (and ligation position for prime-editing samples) was excluded. The function classifyReads(…) was then ran on the mutation table to assign every read to one of the following labels: “pure” if only the expected edit was present; “impure” if the expected edit and some indel(s) were present; “indel” if the expected edit was not present (including if the edit position was deleted) and some indel(s) were present; “scaffold” if nucleotides likely templated from the pegRNA scaffold were present directly after the ligation position; or “reference” if no other label was called. The scaffold label trumped all others. For example, scaffold reads usually included the expected edit but were labelled scaffold, not pure or impure. Detection of scaffold-templated nucleotides is complex as one needs to consider both substitutions and insertions (Supplementary Fig. 2c), different scaffold sequences, and whether the reference happens to match the expected scaffold-templated nucleotides after the ligation position. Details about the detection can be found in the GitHub documentation. To avoid calling sequencing errors, substitutions that were not the expected edit or likely scaffold-templated were ignored in the classification; that is, a “reference” read may include substitutions. Figures like Fig. 1i plots, for each sample, the proportion of reads assigned to each label.

### Indel positions

This refers to Fig. 2c and Supplementary Fig. 7b. Scaffold insertions were excluded from the analysis. To simplify the frequency calculations, we also excluded any read that had more than one indel, which were 311/3028 (10%) for *slc45a2* and 1836/38,553 (5%) for *ctnnb1*. To avoid extrapolating frequencies from too few reads, we excluded any sample with fewer than 50 reads with an indel, this excluded 1/10 samples for *slc45a2* and 1/49 samples for *ctnnb1*. We first computed the frequency of each indel within each sample (number of reads with this indel/total number of reads with an indel), then computed the average frequency of each unique indel across every sample (animal) from each locus, returning one frequency for each indel at each locus. If a sample did not have a given indel, we added a frequency = 0 for this indel in the average frequency calculation. For example, if indel X was only present as 5% of indel reads in one of the nine *slc45a2* samples, we would calculate the average frequency of indel X as the mean of 0.05, 0, 0, 0, 0, 0, 0, 0, 0 (= 0.0056). This calculation sets the total frequency in each plot to be 1.0. Each indel was plotted at a single position: for deletions this was the most 5′ deleted nucleotide; insertions are set by CUTTER as having the same start and stop position.

### Frequencies of deletion lengths after SpCas9 or Cas9 nickase activity

This refers to Supplementary Fig. 12a. We calculated the average frequency of each deletion as above (see *Indel positions*). Briefly, we excluded any deletion that was the desired edit (*cacng2b* 3-bp deletion, or *cacng2b* 6-bp deletion, or *adgrf3b* 3-bp deletion, see Supplementary Fig. 15), only kept reads with a single deletion, and excluded samples with ≤50 deletion reads. We then calculated the frequency of each deletion within each sample (number of reads with this deletion/total number of deletion reads), then the average frequency of each unique deletion across every sample from each locus and condition (SpCas9, D10A, H840A, or PEmax), adding frequency = 0 every time a unique deletion was not observed in a given sample. In the plot, deletion frequencies are summed in 2-bp bins. Each X axis tick corresponds to the last included length of the previous bin. For example, a 2o-bp deletion is counted in the bin left of the 20-bp tick.

### Detection of deletions flanked by microhomologies

The general mutation pattern to be detected was *right microhomology—inner sequence—left microhomology* in the reference sequence with the deleted sequence in the aligned read being *right microhomology—inner sequence* (i.e., *left microhomology* retained) or *inner sequence—left microhomology* (i.e., *right microhomology* retained). 1-bp microhomologies were accepted but the algorithm was designed to always return the longest possible microhomologies. The detection algorithm tested both options (left or right microhomology retained). In occasional cases where both options returned a possible microhomology, the algorithm returned the longest (i.e., most convincing) microhomology. In rare cases where both options gave the same microhomology length, the algorithm returned the right microhomology retained option. Note that right or left microhomology retained may be an arbitrary decision by the alignment algorithm rather than reflect the DNA repair mechanism.

A deletion may be flanked by microhomologies by chance, rather than reflecting repair by MMEJ. This is especially the case as we accepted 1-bp microhomologies: in a random sequence (0.25 chance of each nucleotide), 50% of deletions would technically be flanked by 1-bp microhomologies according to our definition; that is, 0.25 chance that the last nucleotide before the deletion matches the last nucleotide of the deleted sequence (left microhomology retained) + 0.25 chance that the first nucleotide of the deleted sequence matches the first nucleotide after the deletion (right microhomology retained). In practice, the expected probabilities of observing microhomologies by chance will vary by locus. For example, deletions in a highly repetitive sequence will more frequently be flanked by chance microhomologies. To estimate these null probabilities, we simulated 1000 reads with a deletion for each locus. To simulate each deletion read, we first drew a length from a distribution of deletion lengths that is typical from repair of Cas9 DSBs in zebrafish embryos (Supplementary Fig. 19a; see *Fitting distributions of deletion and insertion lengths*). If the length was 3 bp or longer, we considered every deletion of this length that removed the nucleotide that is just 5′ of the expected cut position and drew one deletion at random. If the length was 1 or 2 bp, we considered every deletion of this length that removed at least a nucleotide that is within 4 bp of the expected cut position (i.e., at most 3 bp separating a deleted nucleotide from the nucleotide 5′ of the expected cut position) and drew one deletion at random. We then detected microhomologies from these 1000 simulated reads carrying a deletion as above.

### Detection of insertions templated from neighbouring sequences

Polθ-mediated insertions are often a patchwork of sequences templated from neighbouring sequences (e.g., Supplementary Fig. 17); therefore, naively searching for the entire newly synthesised sequence in neighbouring sequences would rarely map the origin of the inserted sequence. Rather, we aimed for the analysis to extract the longest continuous match between the newly synthesised sequence (insertion and any accompanying substitution(s)) and its neighbouring sequences. To define the neighbouring sequences, we centred the search window on the centre of the newly synthesised sequence. Insertions are set by CUTTER to have the same start and stop position, but an accompanying substitution could shift this centre position. The search window on the reference sequence was defined as this centre position ± 20 bp, then this window was extended further on both sides by the length of the newly synthesised sequence. For example, if the newly synthesised sequence was 10 bp centred on position #100, the search window was defined as the 61-bp sequence from position #70 to #130 (10 bp—20 bp—centre position—20 bp—10 bp). We then detected the longest common substring between the newly synthesised sequence and this search window using function pmatchPattern from R package Biostrings. Polθ can also mediate insertions in reverse-complement orientation ^27^ so we also searched for the longest common substring between the newly synthesised sequence and the reverse-complement of the search window. From the longest common substring in either direction, the algorithm kept the longest one then returned this substring if it was ≥6 bp. We chose this 6-bp minimum length threshold because it brought down false-positive detections in the simulated sample below 25%. This analysis only returned the longest continuous match ≥6 bp for each insertion; therefore, other insertions or parts of an insertion may also be templated but may not be detected as such. Similarly, the analysis was not designed to detect every Polθ-mediated insertion. Polθ can also perform non-templated DNA synthesis^28^ or use sequences far away (>250 bp) from the cut as template ^68^.

As for the deletions, an insertion may match the neighbouring sequences by chance, and not because these neighbouring sequences were genuinely used as template during DNA repair. To estimate these null probabilities, we simulated 1000 newly synthesised sequences for each locus. To simulate each newly synthesised sequence, we first drew a length from a distribution of newly synthesised sequence lengths that is typical from repair of SpCas9 DSBs in zebrafish embryos (Supplementary Fig. 19b; see *Fitting distributions of deletion and insertion lengths*) and generated a random sequence (0.25 chance of each nucleotide) of this length. As for real insertions, we then detected whether ≥6 bp of this random sequence matched neighbouring sequences, centring the search window on the cut position.

### Fitting distributions of deletion and insertion lengths

To fit deletion and insertion lengths that are typical for repair of SpCas9 DSBs in zebrafish embryos (Supplementary Fig. 19a,b), we collected sequencing reads from our previous work ^1,69^ totalling n = 191 embryos mutated across 38 loci using SpCas9 protein complexed with a synthetic crRNA/tracrRNA duplex, for a total of 300,740 sequencing reads. Individual embryos were mutated at 1–9 loci concomitantly but the amplicons were not sufficiently long to detect mutations spanning multiple DSB sites. Reads were aligned with CRISPResso2 and mutations were called with CUTTER (see *Sequencing analysis of prime-editing samples*).

For deletions, we only kept reads carrying some deletion(s) (200,065 reads), then excluded reads carrying more than one deletion, which were 3218/200,065 (2%). To avoid extrapolating frequencies from too few reads with a deletion, we excluded any sample with <100 reads with a deletion, leaving 155 samples across 35 loci. We then calculated, within each sample, the frequency of each unique deletion (number of reads with the deletion/total number of reads with a deletion), then computed the average frequency of each unique deletion across every sample (animal) from each locus, returning one frequency for each indel at each locus. If a sample did not have a given deletion, we added a frequency = 0 in the calculation. For example, if deletion X was only present as 5% of deletion reads in one of six samples from a given locus, we would calculate the average frequency of deletion X as the mean of 0.05, 0, 0, 0, 0, 0 (= 0.008). We then summed, for each locus, the frequencies of all deletions of the same length (e.g., sum of all 5-bp deletion frequencies at locus i), returning one frequency per deletion length at each locus. Finally, we computed the average frequency across loci of each deletion length, adding frequency = 0 each time a length was not found at a given locus. This estimated more general frequencies of each deletion length after repair of Cas9 DSBs in zebrafish embryos (e.g., overall frequency of 5-bp deletions). Supplementary Fig. 19a plots these frequencies, with sum of all frequencies = 1.0. The minimum deletion length was 1 bp, the maximum 74 bp, and the mode 3 bp. We tested various distributions offered by fitdistr from R package MASS to fit this distribution of deletion length frequencies. We chose a log-normal distribution based on visual assessment (Supplementary Fig. 19a) and because it gave a lower (i.e., better fit) D statistic than other distributions in a Kolmogorov–Smirnov test.

For insertions, we only kept reads carrying some insertion(s) (34,489 reads). If a read carried more than one insertion, we only considered one of the insertions, thus keeping the total number of insertion reads the same. To avoid extrapolating frequencies from too few reads with an insertion, we excluded any sample with <50 reads with an insertion, leaving 113 samples across 33 loci. Insertions in zebrafish often appear concomitantly with a multi-bp substitution (see example in Fig. 2e). For each insertion, we included any nearby substitution in the “newly synthesised sequence”. Precisely, any substitution neighbouring the insertion was included in the newly synthesised sequence if it was separated from the insertion by ≤5 bp matching the reference (i.e., a substitution that was >5 bp away from the insertion was not included in the newly synthesised sequence). The following refers to the length of this newly synthesised sequence, rather than the length of the insertion directly. We calculated the average frequency of each length across samples and loci as for the deletions (see above). Supplementary Fig. 19b plots these frequencies, with sum of all frequencies = 1.0. The minimum newly synthesised sequence length was 1 bp, the maximum 95 bp, and the mode 1 bp. We tested various distributions offered by fitdistr from R package MASS to fit this distribution of length frequencies and chose an exponential distribution as it allowed to assign the highest frequency to the shortest length.

### Machine learning pegRNA design tools

We collected edit rates for n = 39 pegRNAs tested in a PE2 RNP encoding 31 edits across 19 genomic loci: 32 pegRNAs from Petri et al., 2022 ^5^; 6 from Vanhooydonck et al., 2025 ^6^; and one from this study (*slc45a2* AA>GG).

For the sequencing data from Petri et al., 2022 ^5^, fastq files were downloaded from NCBI Sequence Read Archive (SRA), BioProject #PRJNA713914 using fastq-dump from SRA Toolkit in command line. Reads were then processed and analysed using CRISPResso2 and CUTTER as for our sequencing data (see *Sequencing analysis of prime-editing samples*). PE3 and PE3b samples were excluded from our analyses. To test machine learning pegRNA design tools (see below), we calculated each sample’s edit rate as the sum of pure and impure edit proportions. To test the pegRNA design tools (Supplementary Fig. 6c), we averaged edit rates from all samples testing the same pegRNA, even if some experimental conditions varied (e.g., synthetic vs. *in-vitro* transcribed pegRNA; or incubation temperature after injection). Metadata were obtained from the authors’ Supplementary Table 6 and 11.

Edit rates from Vanhooydonck et al., 2025 ^6^ were obtained from the authors’ Supplementary Table 1. pegRNA sequences were obtained from Supplementary Table 5.

DeepPrime ^21^ offers 19 models, we excluded models trained for epegRNAs or NRCH PAM, leaving the 9 models listed in Supplementary Fig. 6c. DeepPrime only predicts pegRNAs for edits that are ≤3 bp. This left all pegRNAs encoding a substitution and one pegRNA from Petri et al., 2022 encoding a 3-bp insertion (*adgrf3b* 3-bp insertion), which we excluded as short insertions/deletions generally tend to give higher edit rates than substitutions in zebrafish, leaving n = 27 substitution-encoding pegRNAs to test the DeepPrime models. We obtained DeepPrime predictions for these pegRNAs using the.predict function from genet v0.17.1 Python library. PRIDICT v1 and v2 ^24,25^ pegRNA predictions were obtained using the website pridict.it/. OptiPrime ^23^ pegRNA predictions were obtained using the website optipri.me/. PRIDICT and OptiPrime do not impose an edit size limit but OptiPrime did not generate two of the pegRNAs used by Vanhooydonck et al. 2025, so the correlations were calculated on the complete dataset of n = 39 pegRNAs for PRIDICT and on n = 37 pegRNAs for OptiPrime.

### Base editing sequencing data from Rosello et al., 2022

Sequencing data from Rosello et al., 2022 ^9^ were downloaded as fastq files from NCBI SRA, BioProject #PRJNA825759 using fastq-dump from SRA Toolkit in command line. Metadata were from the authors’ Supplementary Table 10. fastq files were processed and analysed using CUTTER as for our sequencing data (see *Sequencing analysis of prime-editing samples*), but without detection of scaffold incorporations. Samples plotted in Supplementary Fig. 11 are *tyr_5_50k.fastq* (Run #SRR18733203), *tyr_6_50k.fastq* (#SRR18733202), and *tyr_7_50k.fastq* (#SRR18733253).

### Prime editing in mouse embryos

Sequencing data from Kim-Yip et al., 2024 ^29^ were downloaded as fastq files from NCBI SRA, BioProject #PRJNA1040158 using fastq-dump from SRA Toolkit in command line. fastq files were processed and analysed using CUTTER as for our sequencing data (see *Sequencing analysis of prime-editing samples*). We considered all samples injected with PE2 or PEmax mRNA then focused on *Chd2* G>A and *Col12a1* A>C samples as there were more indels to analyse. For each locus, three uninjected control samples were selected at random to be analysed.

### Calculation of PBS:genome Tm

In Supplementary Fig. 6a, we used the R package rmelting to calculate the Tm of the PBS:genome duplex as in Ponnienselvan et al., 2023 ^14^. For example, for the 7-nt PBS used in Supplementary Fig. 6b, the command was: melting(sequence=“GGTCGTC”, nucleic.acid.conc=2e-05, hybridisation.type=’dnarna’, Na.conc=1, method.nn =’sug95’). This command and examples were kindly provided to us by the authors.

### *In-vitro* nickase assay

To assess nickase activity of PEmax protein *in vitro*, we adapted the *in-vitro* prime editing experiment described by Anzalone et al., 2019 ^11^. Fluorescently labelled DNA duplexes (sequences in Supplementary File 1) were produced by annealing two 5′-labelled oligonucleotides (IDT) corresponding to each of the complementary strand for the non-nicked substrate or three oligonucleotides for the pre-nicked substrate. RNPs were made of PEmax protein and gRNA (crRNA/tracrRNA duplex, IDT) complexed at a final concentration of 5 µM and incubated for 10 min at room temperature in 1× reaction buffer (20 mM HEPES pH 7.5; 100 mM KCl; 5% glycerol; 0.2 mM EDTA pH 8.0; 3 mM MgCl2; 5 mM DTT). Next, duplex solution (0.4 µM final concentration) was added to the RNP solution. The reaction was incubated at 37°C for 1 hr. After successive RNase A and proteinase K treatments, the reaction was diluted in a formamide loading buffer (90% formamide, 10% glycerol, 0.01% bromophenol blue). The diluted reaction products were denatured at 95°C before being loaded into a denaturing urea polyacrylamide gel (8 M urea, 1× TBE, 15% polyacrylamide). The products were imaged using the BioRad ChemiDoc imaging system.

### Acridine assay for detection of apoptotic cells

To stain apoptotic cells, we adapted the acridine orange staining assay described by Carrara et al., 2025^7^. In the control experiment (Supplementary Fig. 18), wild-type embryos were collected and incubated at 28.5°C. At 1 dpf, embryos were exposed to one of the following conditions: AOH1160 (MedChemExpress, 20 µM final concentration); CdCl_2_ (Sigma-Aldrich, 20 µM final concentration); or 312–365 nm UV radiation. Note that cadmium (CdCl_2_) is a toxic heavy metal, appropriate safety precautions must be put in place when resuspending the powder. For UV exposure, embryos were transferred to glass beakers and placed on a UV transilluminator used to cut bands from agarose gels (ECX-F20.SkyLight). UV was set to 100% and embryo were irradiated for 30 min. All embryos were then returned to the incubator. AOH1160 and CdCl_2_ were washed-out the next morning (2 dpf, ∼24 hr treatment) before acridine orange staining. In the *polq* mutant experiment (Fig. 4c), wild-type and *polq* mutant embryos were injected at the one-cell stage with PEmax RNPs carrying a targeting pegRNA (*slc45a2* GG>AA) or a pegRNA targeting a *BFP* transgene not present in these animals (sequence in Supplementary File 1), which served as non-targeting control for the effect of RNP injections. Acridine orange staining (see below) was then performed the next day at 1 dpf.

At 1 dpf (experiment with *polq* mutants) or 2 dpf (control experiment), embryos were pooled in groups of 20 and incubated in a 3 µg/mL acridine orange solution (Sigma-Aldrich) in egg medium for 30 min in the dark at room temperature. After staining, embryos were washed repeatedly in egg medium (without methylene blue) using a mini see-saw rocking shaker (100 oscillations/min), then transferred to fresh egg medium in a clean Petri dish. To facilitate imaging, embryos were manually dechorionated using fine forceps. Fluorescence imaging was performed using a Leica MSV269 microscope mounted with a Leica DFC310 FX camera. In the LAS X software, acquisition was set to black-and-white fluorescence mode. Acridine-positive cells exhibited green fluorescence (emission peak at 525 nm) when excited with a blue filter (excitation peak at 502 nm). The yolk tended to fluoresce brightly even after repeated wash-outs so we laid flat each larva as best as possible to focus the photograph on the tail. Images were analysed using Fiji v2.14.0 ^70^. For each image, we segmented the tail region from the cloaca to the tail fin, set a minimum fluorescence intensity threshold to exclude background signal, and counted fluorescent puncta using Analyze > Analyze particles.

Images in Fig. 4c were acquired with an Olympus FV1000 laser scanning confocal microscope using a 20× 1.0 NA objective lens (Zeiss, W Plan-Apochromat 20×/1.0). Z-volumes were captured at a z-plane spacing of 3 µm; resolution was 1024 × 1024 pixels with a scan rate of 4 µs/pixel.

### Pictures

For the pictures in Fig. 3d and Fig. 4d, larvae were anaesthetised with tricaine (MS-222) and pictures were taken with a Leica DFC310 FX camera mounted on a Leica MSV269 brightfield microscope with illumination from below the sample.

### Variant types from ClinVar

This refers to Supplementary Fig. 1. We downloaded file variant_summary.txt.gz from ClinVar’s ftp site in December 2024. We used variants on the GRCh38 (hg38) human genome reference, removed duplicated variants, and only kept variants in protein-coding regions (“p.” in variant name; e.g., “p.Gly262Arg”). The variants we labelled “premature stop” were those labelled as nonsense (“Ter” in variant name; e.g., “p.Tyr47Ter”) or frameshift (“fs” in variant name; e.g., “p.Phe468fs”). Category “other” included the following types: “Indel”, “Inversion”, “Translocation”, “Variation”, “copy number gain”, “copy number loss”, “Duplication”, “Microsatellite”.

### Statistics

Threshold for statistical significance was 0.05. In figures, ns refers to p > 0.05, * to p ≤ 0.05, ** to p ≤ 0.01, *** to p ≤ 0.001. In text, data distributions are reported as mean ± standard deviation, unless stated otherwise.

### Software

Data analysis was performed in R v4.4.2 ran through RStudio 2025.09.2+418; plots were generated with R package ggplot2. Some analyses used Jupyter notebooks written in Python3. Figures were prepared with Adobe Illustrator 2019.

## Data and data availability

Source code of the CUTTER R package is available at github.com/francoiskroll/cutter.

Sequencing data will made available on NCBI SRA.

## Acknowledgements

This work was supported by the Fondation pour la Recherche Médicale grant no. SPF202309017505 awarded to F.K and the European Commission HORIZON-PathFinder EdiGenT grant no. 101070903.

We thank the members of the Del Bene lab for helpful discussions and Karine Duroure and Fish Facility staff for fish husbandry. We thank Alexandra Lubin, Katie-Jo Thorpe, and Tanvi Tambaki (Payne lab, UCL Cancer Institute, London) for the Illumina MiSeq runs; Guillaume Pézeron (Muséum National d’Histoire Naturelle, Paris) for kindly giving access to his microinjector and donating the *polq* KO line; Talya Goble and Jason Rihel (UCL, London) for feedback on the draft manuscript.

---

**Supplementary Fig. 1.**
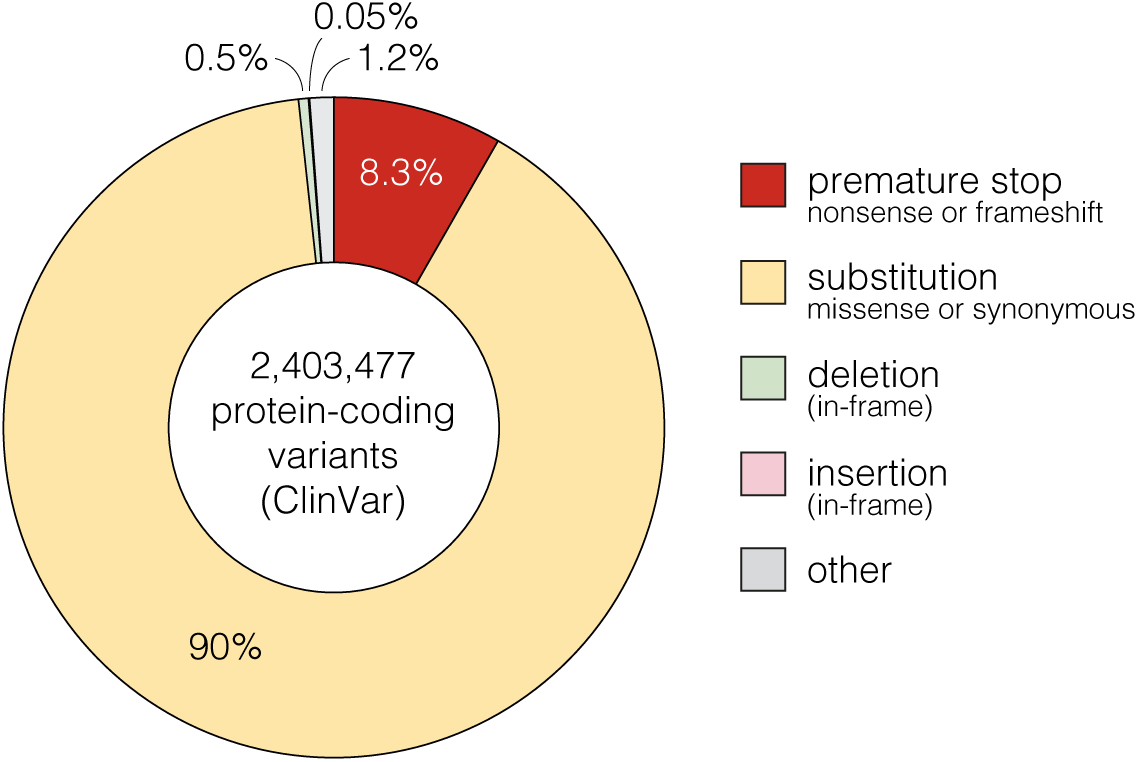
Types of human variants recorded in ClinVar. Proportions of different types of human genomic variants recorded in the ClinVar database. Only variants in protein-coding sequences were included.

**Supplementary Fig. 2.**
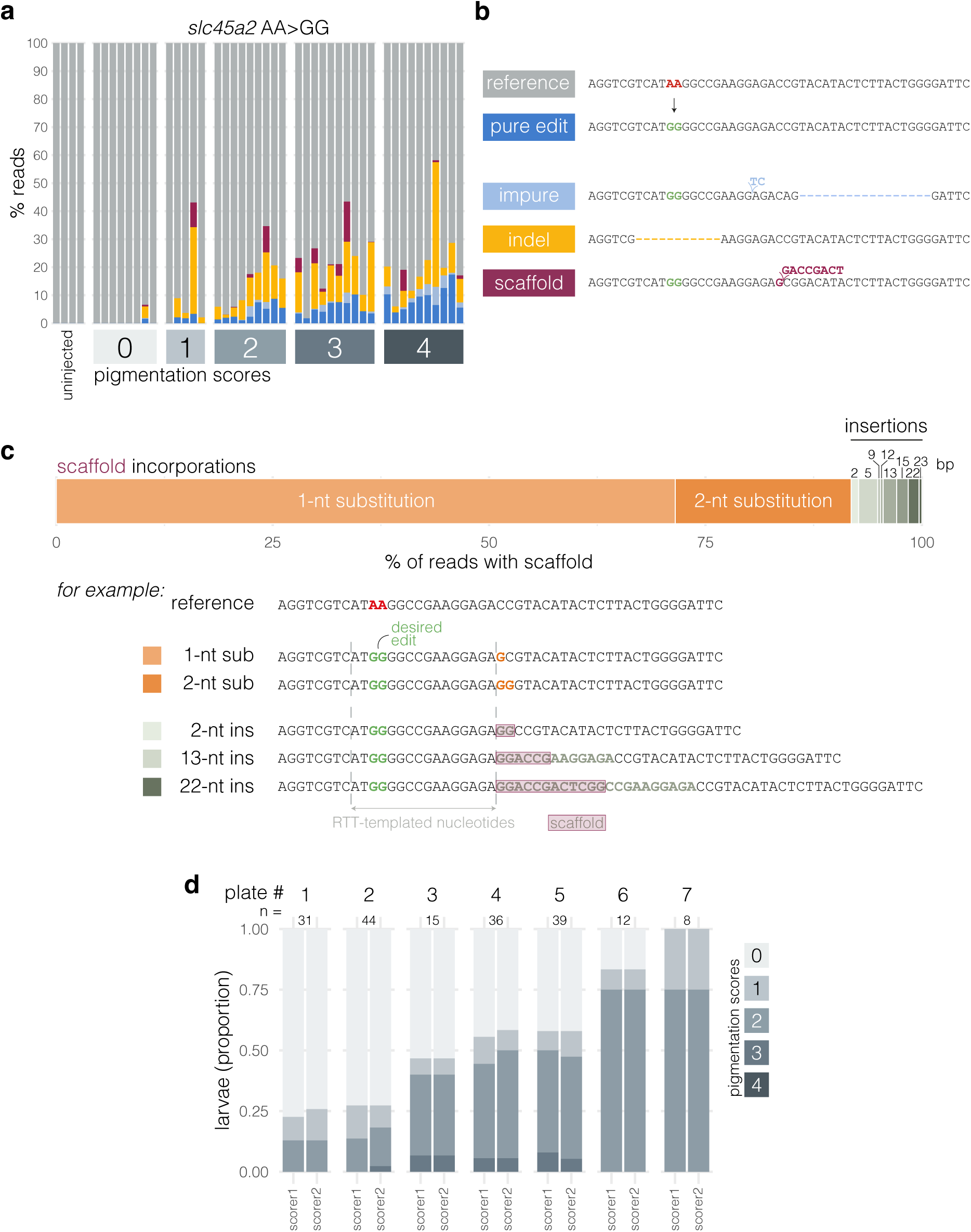
Development of the pigmentation recovery assay for quick readout of prime editing rates. a, Sequencing results from larvae grouped according to their pigmentation score. Each bar represents an individual larva, with colours representing the percentage of reads assigned to each label, see b. Minimum sequencing depth was 131×. n = 42 injected samples. **b,** Examples of sequencing reads. Every read is assigned a single label depending on its mutation calls: “reference” if the read matches the reference sequence; “pure edit” if only the expected edit (AA>GG) is present; “impure edit” if the expected edit and one or more indel(s) are present; “indel” if the read does not carry the edit but one or more indel(s); and “scaffold” if a sequence likely templated from the pegRNA’s scaffold is present. **c,** Scaffold incorporations can include both substitutions and insertions. (top) Of 1274 reads with a scaffold incorporation from the *slc45a2* prime-edited samples in Fig. 1i, 1170 (92%) had a scaffold substitution (orange), 104 (8%) had a scaffold insertion (khaki). (bottom) Examples of reads with a scaffold incorporation. For insertions, the nucleotides that match the scaffold sequence are framed in purple. sub, substitution; ins, insertion. **d,** Concordance in pigmentation scores between two scorers assessing the same injected larvae. Colours in each bar plot represent the proportion of larvae assigned to each pigmentation score.

**Supplementary Fig. 3.**
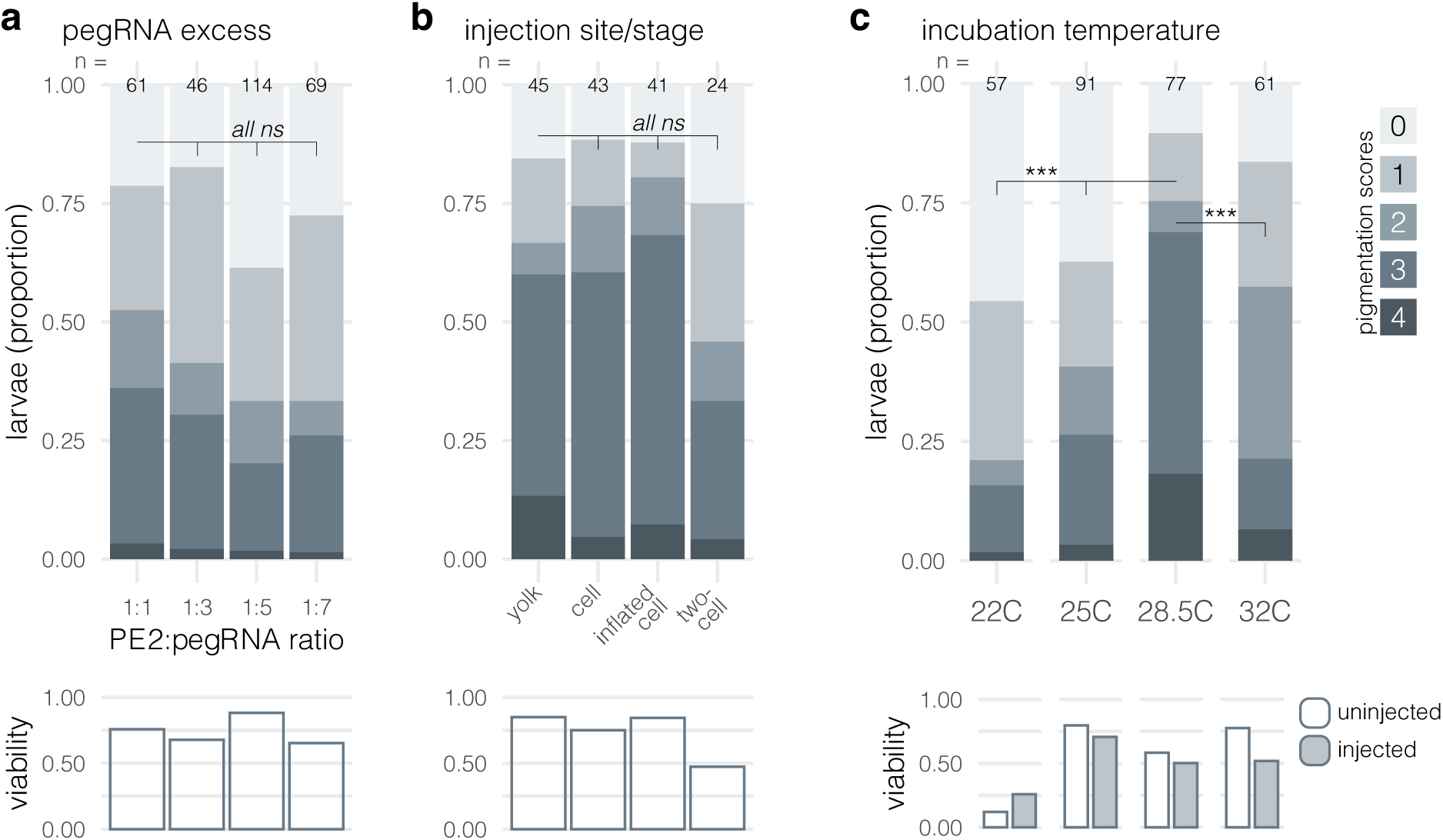
Optimisation of PE RNP injection conditions using the pigmentation recovery assay. a, As the pegRNA PBS is fully complementary to the spacer, the pegRNA may misfold, blocking its activity ^14^. Using an excess of pegRNA could benefit prime editing rates by providing the PE protein with more correctly folded pegRNA. (top) Here, phenotypic penetrance (proportion of larvae assigned to each pigmentation score) when an excess of pegRNA was used. PE2 protein was kept constant at 26 fmol while pegRNA was gradually increased. 1:1: 26 fmol, 1:3: 78 fmol, 1:5: 130 fmol, 1:7: 182 fmol pegRNA. ns p > 0.07 by Fisher’s exact test. (bottom) Viability as proportion of larvae assessed as viable at 6 dpf, normalised to uninjected embryos. **b,** Solution can be injected in the zebrafish egg’s yolk or cell. Yolk injections are quicker but the RNP must first cross into the cell. Cell injection may be beneficial by avoiding this delay. In mouse zygotes, injections at the two-cell stage increased edit rates ^29^. (top) Here, phenotypic penetrance (pigmentation scores) depending on injection site. Embryos were injected with 30 fmol (1 nL) of PE2/pegRNA RNP, except two-cell stage embryos which received 30 fmol (1 nL) of RNP in each cell, so 60 fmol (2 nL) total. ns p > 0.32 by Fisher’s exact test. (bottom) Viability as proportion of larvae assessed as viable at 5 dpf, normalised to uninjected embryos. **c,** The standard incubation temperature for zebrafish embryos is 28.5°C but as ectotherms can develop in a range of temperatures. Cooler incubation temperatures likely benefit Cas9 nuclease mutagenesis by delaying the first divisions ^71^. In contrast, warmer incubation at 32°C increased prime editing rates in a previous zebrafish study ^5^, potentially because this is closer to the optimal temperature of the RT^72^. (top) Here, phenotypic penetrance (pigmentation scores) after incubation at different temperatures. Embryos were incubated for 3 hr directly after injection at the corresponding temperature then transferred to 28.5°C. *** p < 0.001 by Fisher’s exact test. (bottom) Viability as proportion of larvae assessed as viable at 6 dpf. We also tested 18°C but too few embryos survived (6/136).

**Supplementary Fig. 4.**
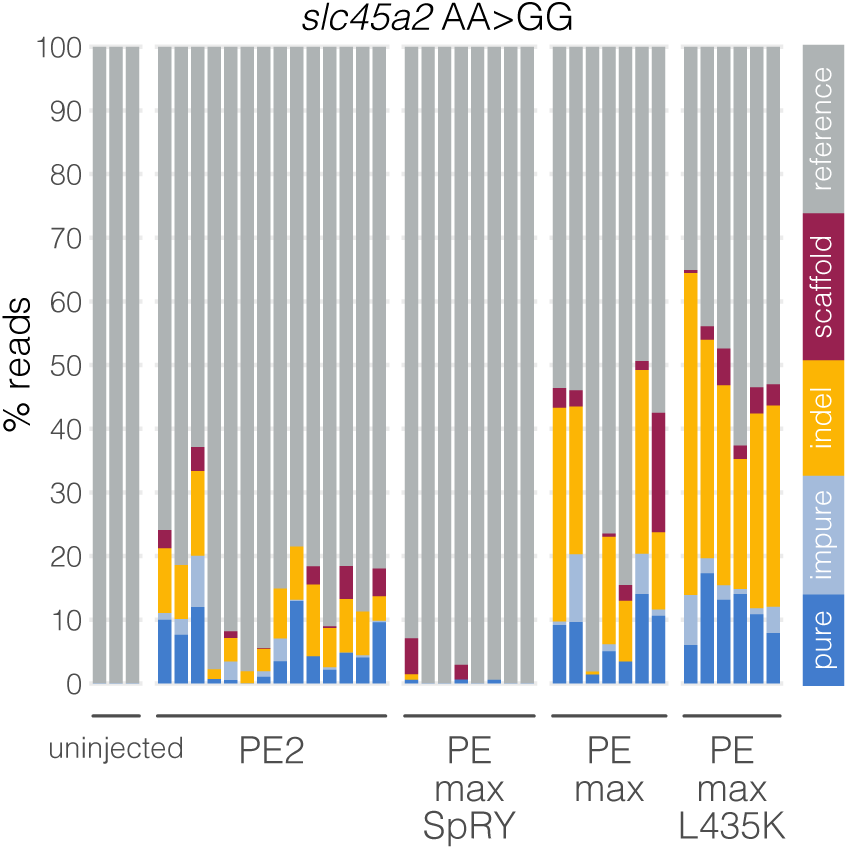
Test of PE protein variants at *slc45a2* AA>GG. Sequencing results from larvae injected with 30 fmol of the corresponding PE RNP. Each bar represents an individual larva, with colours representing the percentage of reads assigned to each label. Minimum sequencing depth was 766×. n = 35 injected samples.

**Supplementary Fig. 5.**
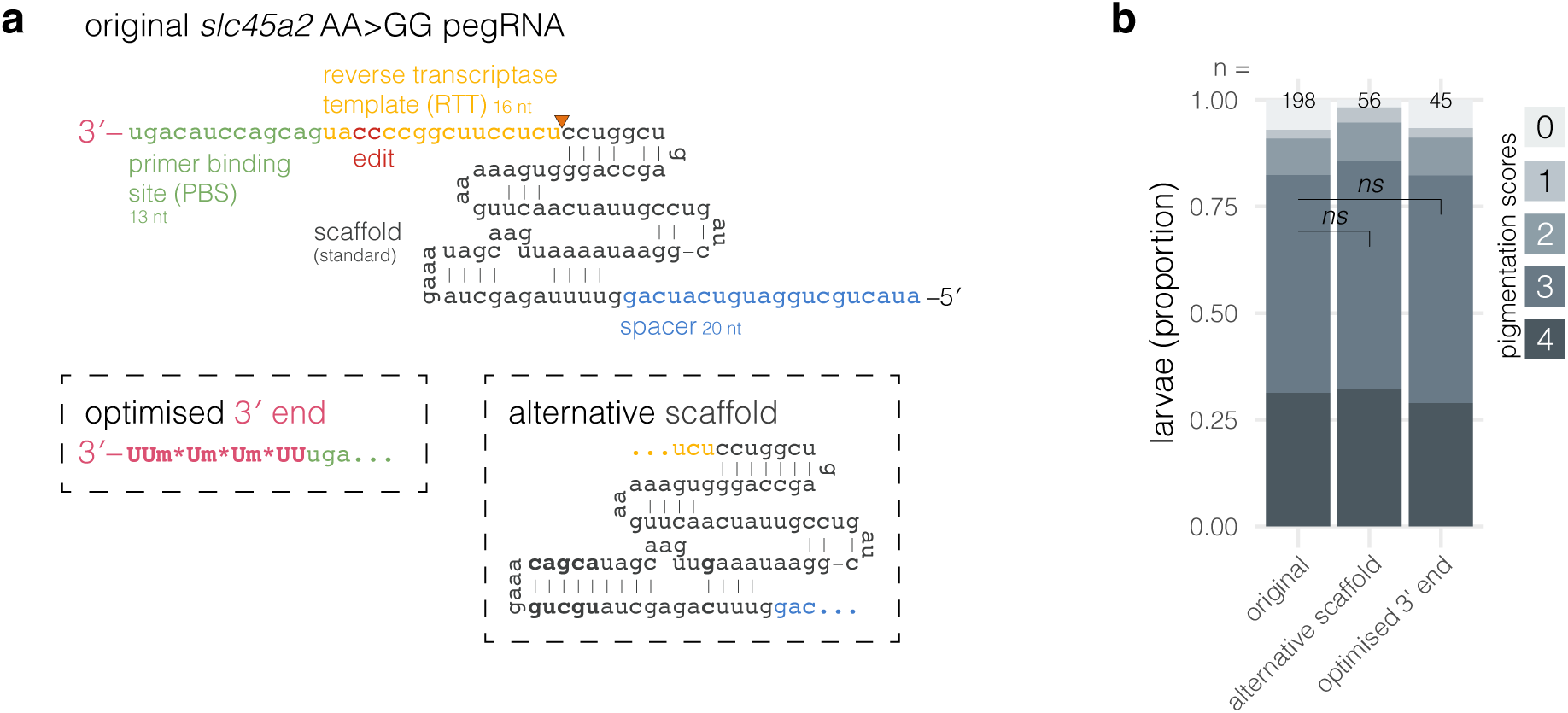
Test of pegRNA modifications at *slc45a2* GG>AA. **a**, The full sequence of the original *slc45a2* pegRNA used in the pigmentation recovery assay. The orange arrowhead indicates the position of the riboabasic spacer tested in Fig. 1i. (bottom left) The chemically modified 3′ end from Yan et al., 2024 ^19^. Um denotes a uridine carrying a 2′-O-methylation; * denotes a phosphorothioate bond. (bottom right) The alternative pegRNA scaffold from Dang et al., 2015 ^20,21^. Compared to the standard scaffold, the tetraloop is extended by 5 nt and a uridine in a four-uridine stretch is substituted to cytidine (nucleotides in bold). **b**, Phenotypic penetrance (proportion of larvae assigned to each pigmentation score) in larvae injected with 30 fmol of PEmax^L435K^ complexed with the original *slc45a2* AA>GG pegRNA, or the alternative scaffold pegRNA, or the optimised 3′-end pegRNA. ns p > 0.6 by Fisher’s exact test.

**Supplementary Fig. 6.**
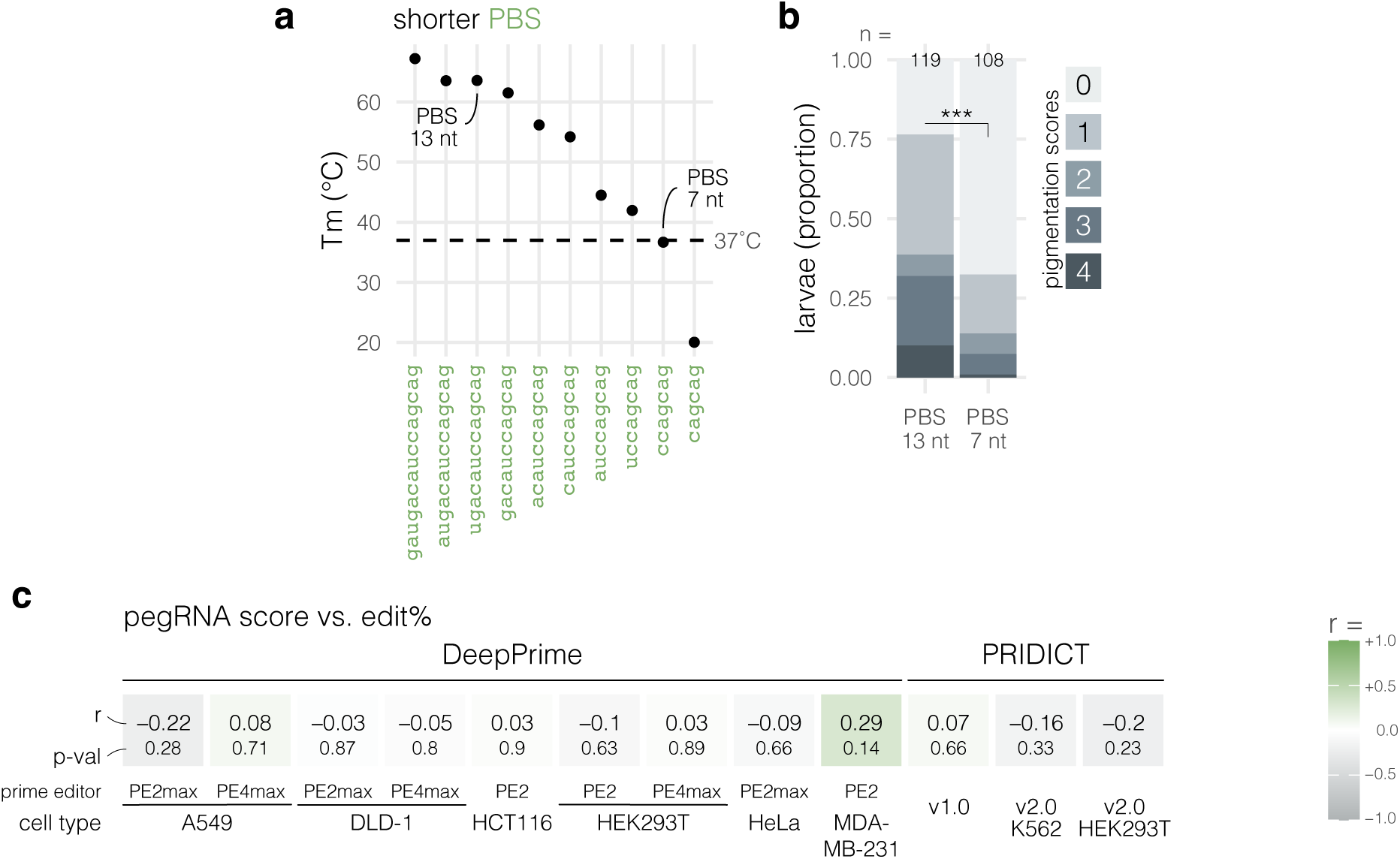
Assessment of pegRNA design tools in zebrafish embryos. a, Tm calculations of the PBS:genome duplex as the PBS of the *slc45a2* AA>GG pegRNA is shortened from 15 to 6 nt. **b,** Phenotypic penetrance (proportion of larvae assigned to each pigmentation score) in larvae injected with 30 fmol of PE2 protein complexed with the original 13-nt PBS *slc45a2* AA>GG pegRNA or its 7-nt PBS counterpart. *** p < 0.001 by Fisher’s exact test. **c,** Predictions of pegRNA editing efficiencies in zebrafish embryos by 13 machine learning models. Tile colours represent Pearson’s correlations (r from −1 to +1). In each tile, the top number is Pearson’s r, the bottom is its p-value. Each correlation is calculated between the pegRNAs’ predicted scores and their average edit rates (sum of pure and impure edit rates) in zebrafish embryos. Median number of samples per pegRNA was 5. For PRIDICT ^24,25^: n = 39 pegRNAs, which is the complete dataset. For OptiPrime ^23^: n = 37 pegRNAs as it did not generate two pegRNAs. DeepPrime ^21^ only works on edits ≤3 bp so not every pegRNA could be included: n = 27 pegRNAs. Including only the substitution-encoding pegRNAs (same set as for DeepPrime) in the PRIDICT and OptiPrime analysis did not change the interpretation (r = −0.28 to −0.08). We also tested the correlation between the pegRNAs’ “unintended scores” predicted by PRIDICT v1 and their average indel rates: r = −0.04, p = 0.8. Data from Petri et al., 2022 ^5^ (n = 32 pegRNAs); Vanhooydonck et al., 2025 ^6^ (n = 6 pegRNAs); and the *slc45a2* AA>GG pegRNA used with PE2 RNP in this study. Embryos were injected with ∼6 fmol (30 fmol for *slc45a2* AA>GG) of RNP made of PE2 protein and *in-vitro* transcribed or synthetic pegRNA.

**Supplementary Fig. 7.**
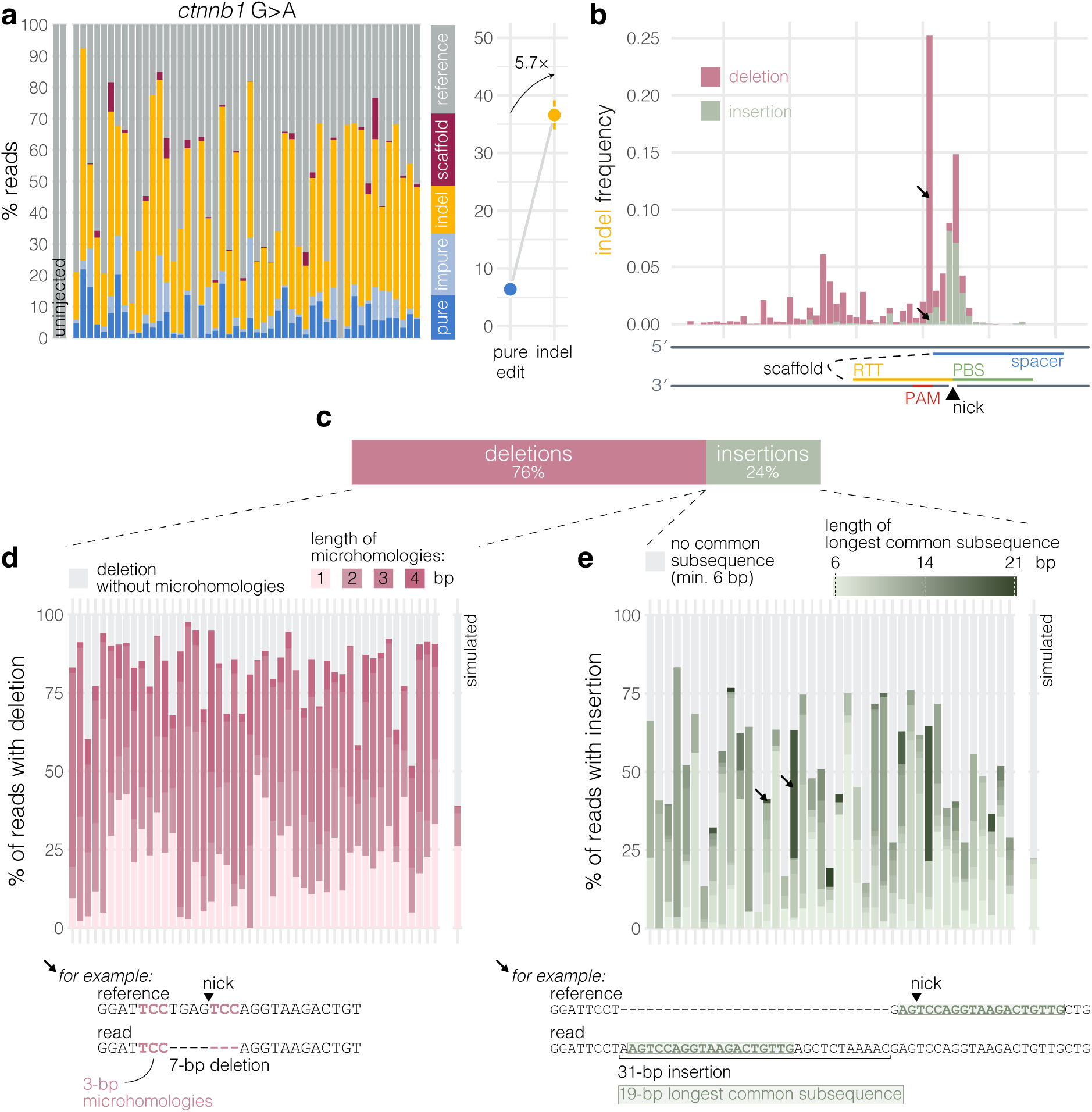
Indels generated during prime editing at *ctnnb1* show signatures of MMEJ. a, (left) Sequencing results from larvae injected with 30 fmol PEmax complexed with pegRNA encoding a one-bp substitution (G>A) in *ctnnb1*. Each bar represents an individual larva, with colours representing the percentage of reads assigned to each label. Minimum sequencing depth was 456×. n = 60 injected samples. (right) The pure edit rate compared to the indel rate (mean ± SEM). **b,** Frequency of prime editing-generated indels across genomic positions at the *ctnnb1* locus, from n = 48 of the samples in a. Each frequency represents the average frequency across larvae of indels starting at this position. Each indel (n = 639 unique indels observed) contributes to the frequency once at its start position, which is the most 5′ nucleotide involved. For example, a 7-bp deletion which deleted nucleotide #71–77 is counted at #71. Scaffold incorporations are not included. **c,** In the average *ctnnb1* prime-edited sample, 76% of reads with an indel had a deletion, the other 24% had an insertion. **d,** (top) Of all sequencing reads with a deletion, percentage which showed microhomologies flanking the deletion. Each bar represents an individual prime-edited larva. Darker pink colours represent longer microhomologies, grey represents deletions without microhomologies. Only samples with ≥50 reads with a deletion were analysed. There were 102–1561 reads with a deletion per sample included. The “simulated” sample represents 1000 simulated deletions. (bottom) Example of a 7-bp deletion flanked by 3-bp microhomologies found in all (48/48) samples plotted; the arrow in b indicates its genomic position. **e,** (top) Of all sequencing reads with an insertion, percentage which had a continuous match ≥6 bp between the “newly synthesised sequence” and the neighbouring sequences. Insertions generated during DSB repair in zebrafish are often accompanied with a multi-bp substitution (see example in Fig. 2e), they together form the “newly synthesised sequence” which is compared to the neighbouring sequences to find a possible match. Each bar represents an individual prime-edited larva. Darker green colours represent longer matches between the newly synthesised sequence and the neighbouring sequences, grey represents newly synthesised sequences with a <6 bp match; that is, whose length was too short to analyse, or were untemplated, or used a template that was further away. Only samples with ≥50 reads with an insertion were analysed. There were 54–1535 reads with an insertion per sample included. The “simulated” sample represents 1000 newly synthesised sequences generated at random. (bottom) Example of a 31-bp insertion with at least 19 bp templated from a neighbouring sequence (green); the arrows in b and e indicate its genomic position and the two samples in which it was found.

**Supplementary Fig. 8.**
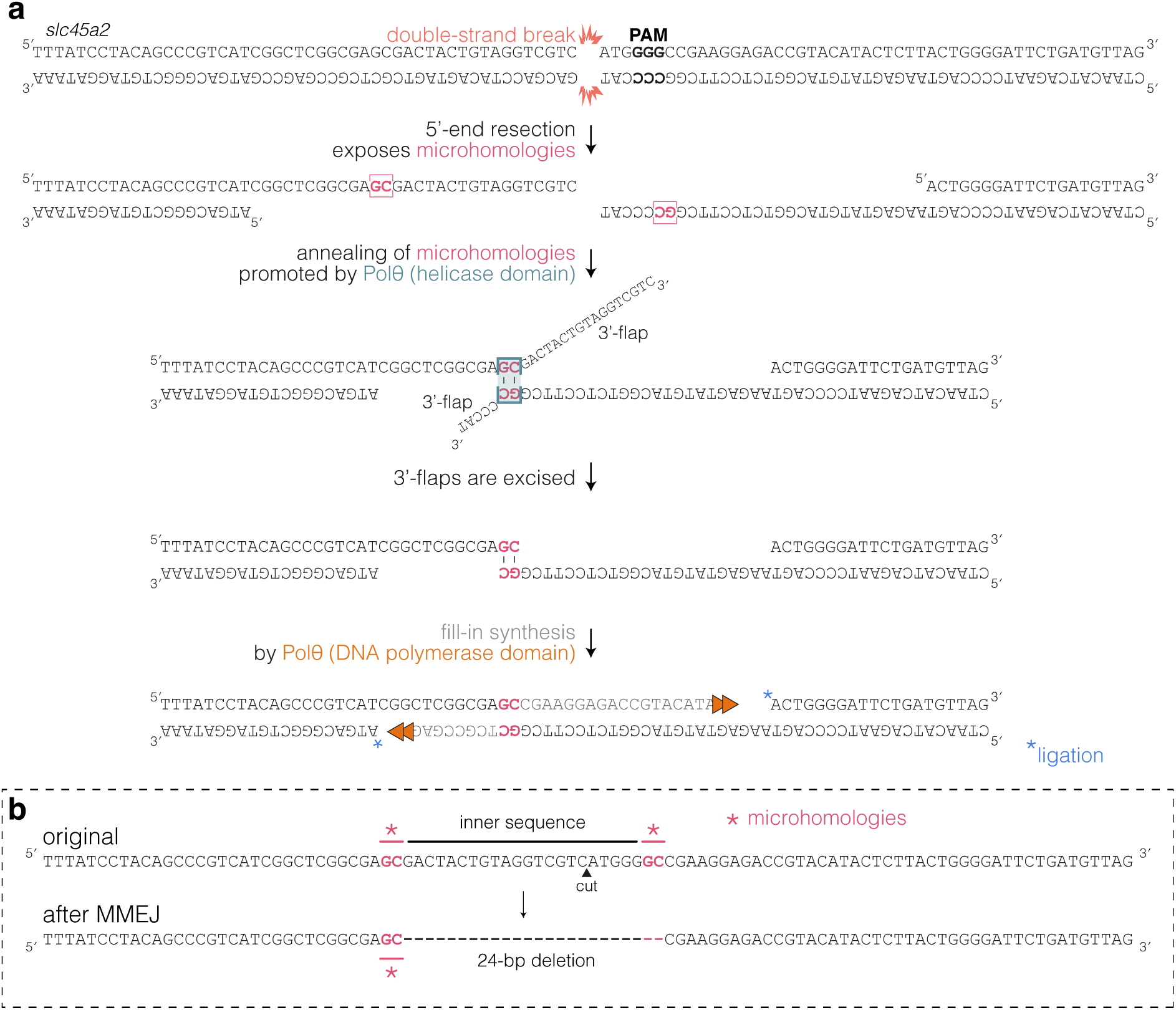
Canonical MMEJ mechanism generating microhomology-flanked deletions. a, As example, simplified MMEJ mechanism creating a 24-bp deletion observed in embryos mutated with SpCas9 at the *slc45a2* locus (0.44—58% in 3/23 samples). **b,** Mutation generated by the MMEJ mechanism in a. The typical MMEJ deletion removes the inner sequence between the two microhomologies and one microhomology. Whether the left or right microhomology is shown as deleted may be an arbitrary decision by the alignment algorithm.

**Supplementary Fig. 9.**
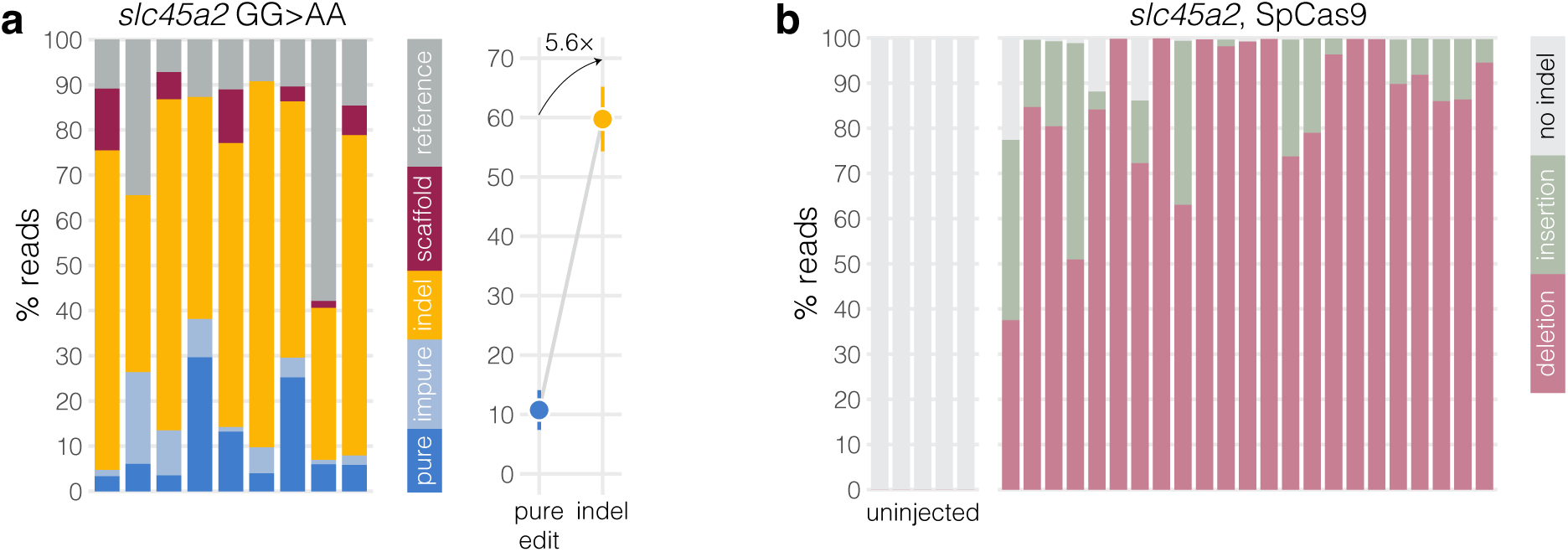
Prime editing and SpCas9 indels at *slc45a2* in wild-type embryos. a, (left) Sequencing results from wild-type larvae injected with 30 fmol PEmax complexed with the *slc45a2* GG>AA pegRNA. Each bar represents an individual larva, with colours representing the percentage of reads assigned to each label. Minimum sequencing depth was 118×. n = 9 samples. The same samples are plotted in Supplementary Fig. 15. (right) The pure edit rate compared to the indel rate (mean ± SEM). **b,** Sequencing results from wild-type larvae injected with 30 fmol SpCas9 complexed with gRNA targeting the same *slc45a2* locus (same spacer sequence) as in a. Each bar represents an individual larva, with colours representing the percentage of reads with an insertion, a deletion, or neither. Per larva, 98 ± 6% of reads carried an indel. Minimum sequencing depth was 788×. n = 23 injected samples.

**Supplementary Fig. 10.**
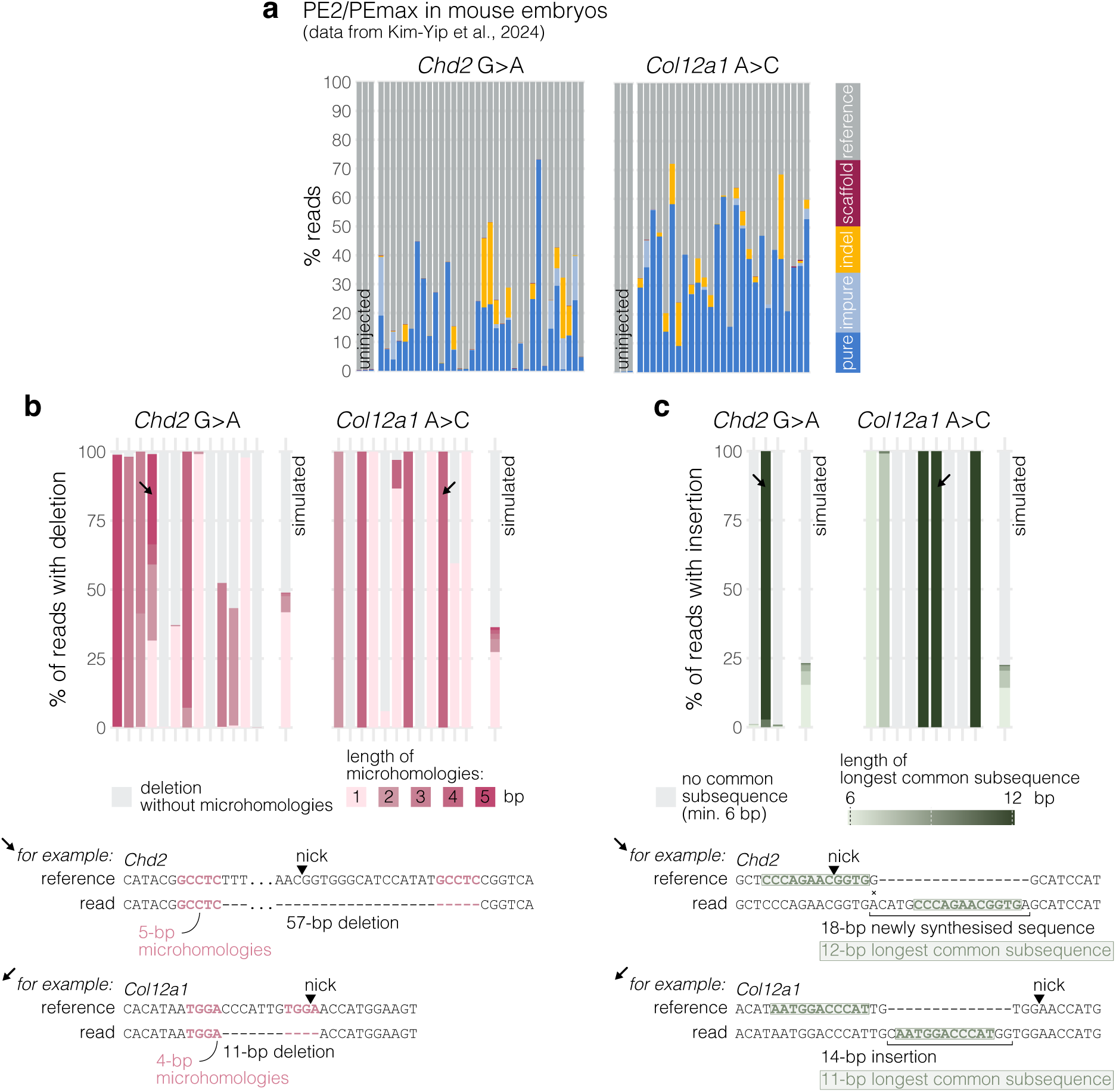
Prime editing causes indels with signature of MMEJ in mouse embryos. a, Sequencing results from mouse embryos injected with synthetic pegRNA and PE2 (*Chd2*) or PEmax (*Col12a1*) mRNA. For *Chd2*, embryos were injected at the zygote or two-cell stage. For *Col12a1*, embryos were injected at the two-cell stage. Each bar represents an individual embryo, with colours representing the percentage of reads assigned to each label. Minimum sequencing depth was 1579×. *Chd2*: n = 34 injected samples; *Col12a1*: n = 27 injected samples. Sequencing data from Kim-Yip et al., 2024 ^29^. **b,** (top) Of all sequencing reads with a deletion, percentage which showed microhomologies flanking the deletion. Each bar represents an individual prime-edited mouse embryo. Darker pink colours represent longer microhomologies, grey represents deletions without microhomologies. Only samples with ≥50 reads with a deletion were analysed. There were 104–4589 reads with a deletion per sample included. The “simulated” samples represent 1000 simulated deletions at each locus. (bottom) Examples of deletions with microhomologies; the arrows indicate the samples in which they were found. **c,** (top) Of all sequencing reads with an insertion, percentage which had a continuous match ≥6 bp between the “newly synthesised sequence” and the neighbouring sequences. Each bar represents an individual prime-edited embryo. Darker green colours represent longer matches between the newly synthesised sequence and the neighbouring sequences, grey represents newly synthesised sequences with a <6 bp match; that is, whose length was too short to analyse, or were untemplated, or used a template that was further away. Only samples with ≥50 reads with a deletion were analysed. There were 64–1368 reads with an insertion per sample included. The “simulated” samples represent 1000 newly synthesised sequences generated at random for each locus. (bottom) Examples of newly synthesised sequences that were templated from a neighbouring sequence (green); the arrows indicate the samples in which they were found.

**Supplementary Fig. 11.**
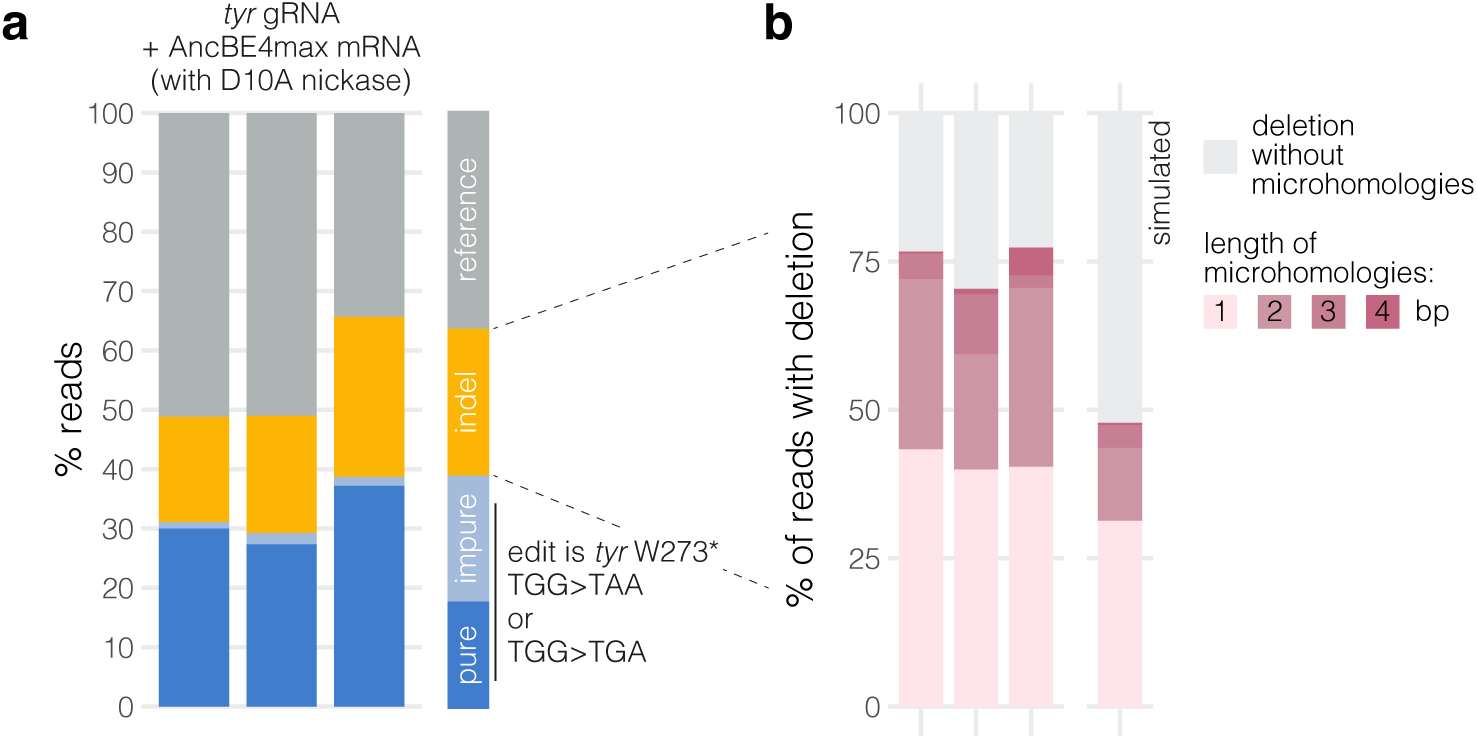
Base editing using the D10A nickase also generates deletions with signature of MMEJ. a, Sequencing results from larvae injected with mRNA encoding the C-to-T base editor AncBE4max and a gRNA targeting *tyr*. Each bar represents sequencing data from a pool of zebrafish larvae, with colours representing the percentage of reads assigned to each label. Minimum sequencing depth was 44,859×. Sequencing data from Rosello et al., 2022 ^9^. **b,** Of all sequencing reads with a deletion, percentage which show microhomologies flanking the deletion. Darker pink colours represent longer microhomologies, grey represents deletions without microhomologies. Only samples with ≥50 reads with a deletion were analysed. There were 7300–12,230 reads with a deletion per sample. The “simulated” sample represents 1000 deletions.

**Supplementary Fig. 12.**
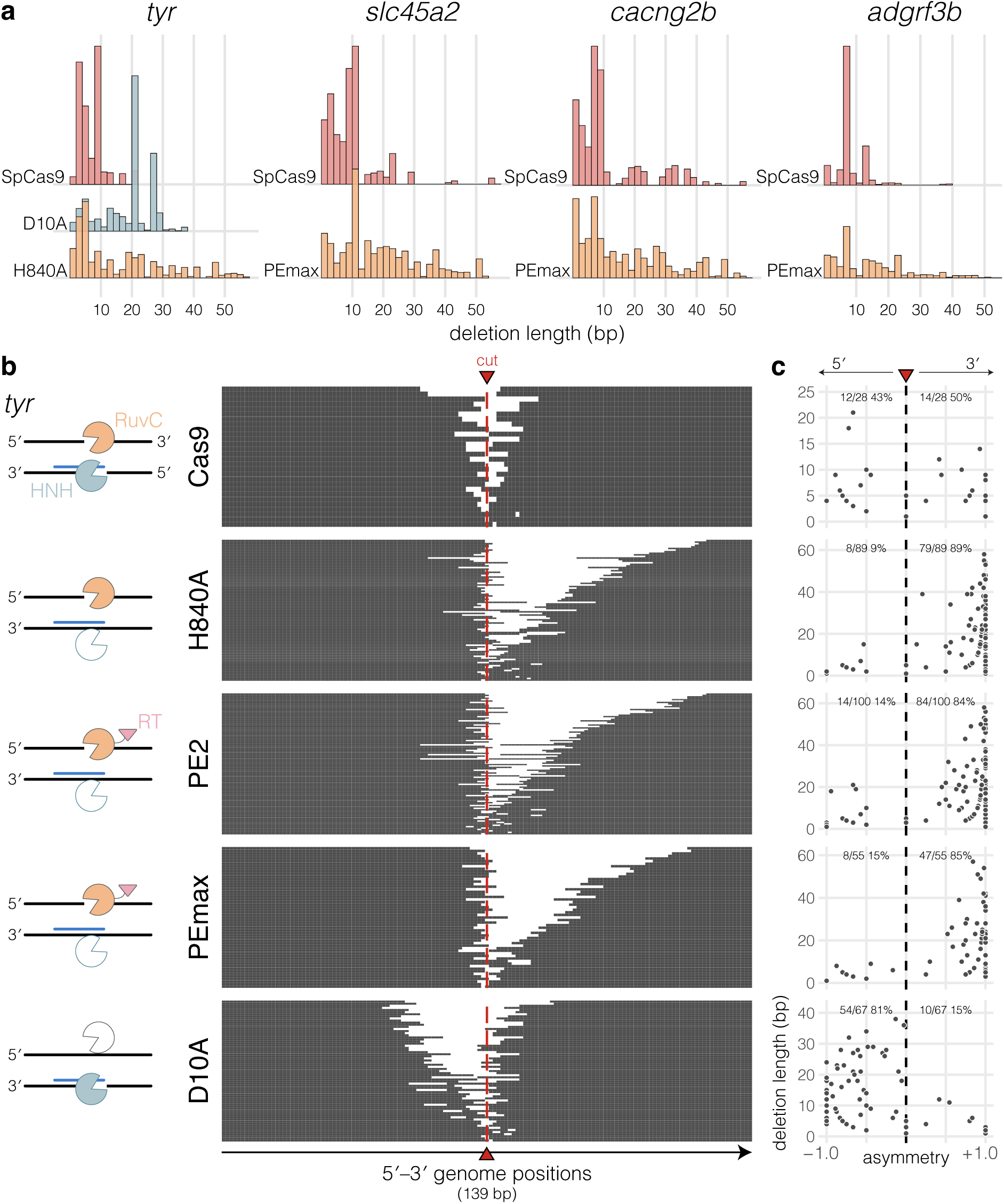
Cas9 nickase activity results in long asymmetric deletions, characteristic of fork-MMEJ of replication-associated single-ended DSBs. a, Frequencies of deletion lengths after SpCas9, D10A nickase, H840A nickase, or PEmax/pegRNA. For *tyr*, H840A also includes deletions found in PE2/gRNA and PEmax/gRNA-injected samples (experiment in Fig. 3b–f). For *slc45a2*, *cacng2b*, and *adgrf3b*, SpCas9 was complexed with a gRNA and PEmax was complexed with a pegRNA carrying the same spacer sequence (wild-type samples in Supplementary Fig. 15). **b,** Deletion patterns at the *tyr* locus when both genomic strands were cleaved by SpCas9; or when the non-target strand was cleaved by H840A nickase, PE2, PEmax; or when the target strand was cleaved by D10A nickase. Each protein was complexed with a gRNA. Each row is a unique deletion allele aligned to the reference amplicon, plotted from the shortest deletion at the top to the longest at the bottom. Nucleotides present are in grey, white gaps are deleted nucleotides. The red line indicates the predicted cut position. The cartoons on the left indicate which genomic strand was cleaved in each condition. Sequencing data from the experiment in Fig. 3b–f. **c,** Deletion length (bp) as a function of deletion asymmetry. Each dot is a unique deletion allele. Asymmetry is the ratio of deleted nucleotides on either side of the cut. A deletion that is entirely 5′ of the cut position has asymmetry = −1.0; a deletion that is entirely 3′ of the cut position has asymmetry = +1.0; a deletion with asymmetry = 0 deleted exactly the same number of bp on either side of the cut.

**Supplementary Fig. 13.**
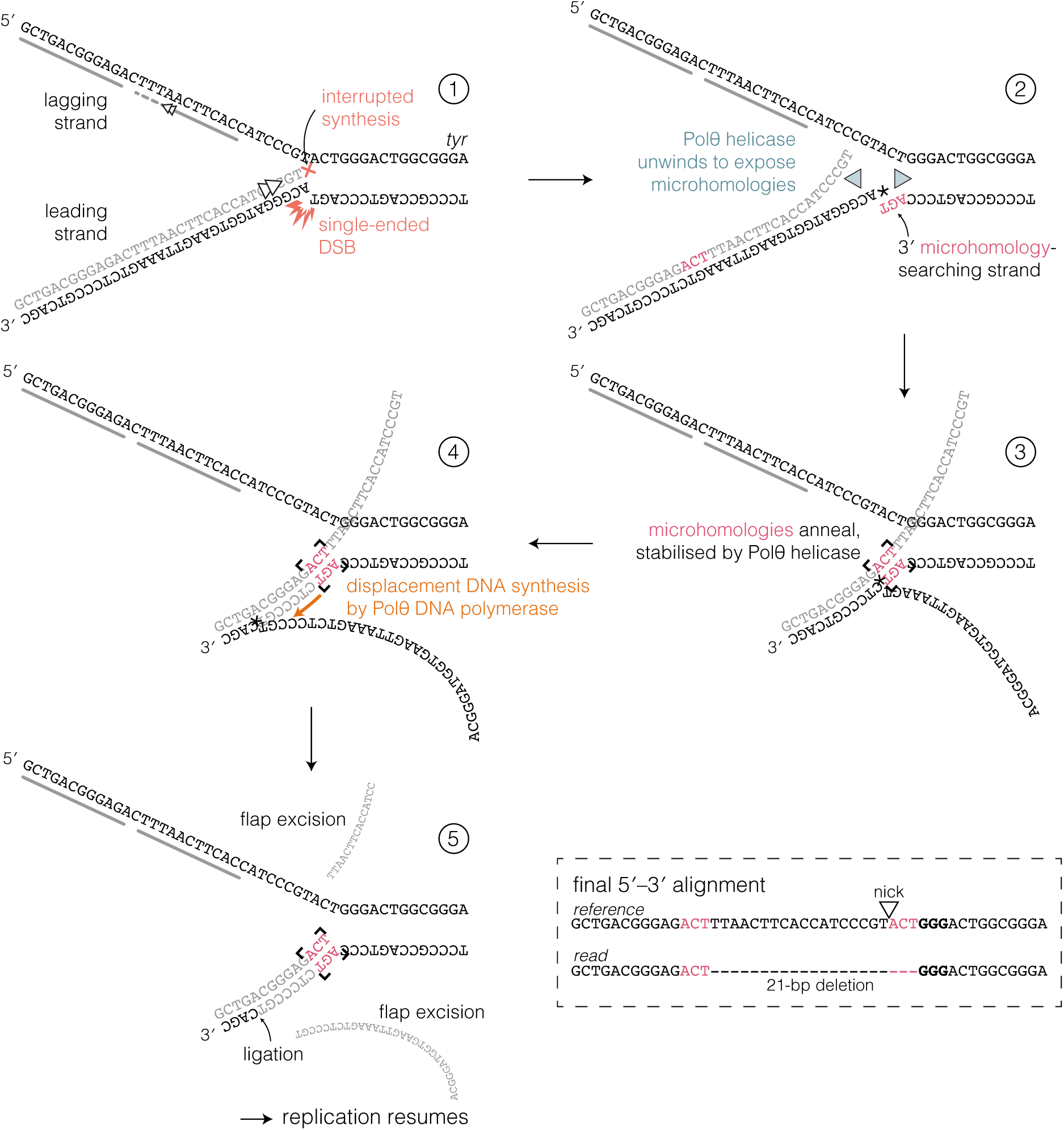
Potential repair mechanism of a replication-associated single-ended DSB by fork-MMEJ. Fork-MMEJ mechanism as proposed by Li et al. (2026) ^41^, here taking as example the most frequent deletion found in larvae injected with D10A nickase targeting *tyr* (average 34% of deletion reads in n = 10 samples, see peak in Supplementary Fig. 12a). Asterisk (*) in 2–4 indicates the nicked position. The fork-MMEJ mechanism varies depending on the direction of replication and whether the leading or lagging strand is cleaved. Here, we assume the depicted configuration. (framed) Deletion generated by the fork-MMEJ mechanism, aligned to the reference.

**Supplementary Fig. 14.**
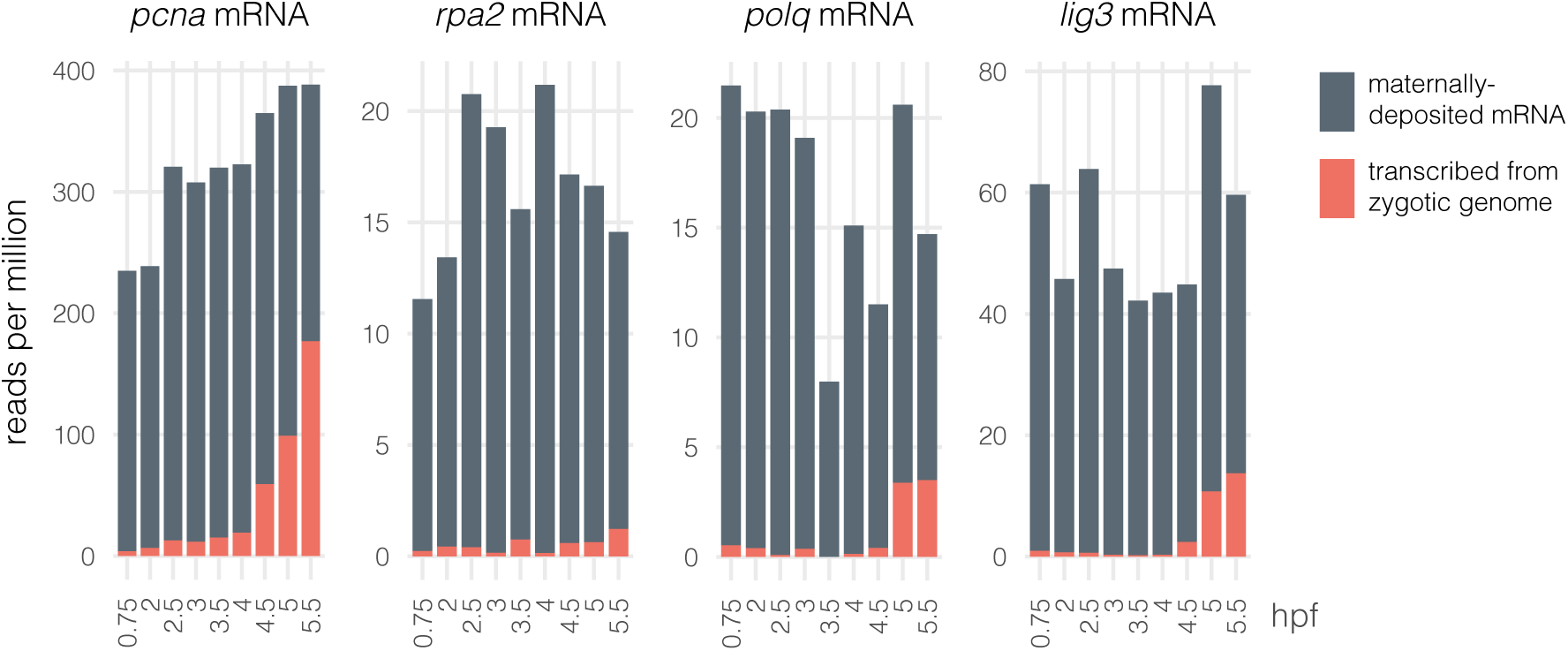
The main MMEJ and fork-MMEJ factors are maternally deposited. Transcript levels of the core MMEJ/fork-MMEJ factors ^41^ during early zebrafish embryogenesis (0.75–5.5 hours post-fertilisation, hpf) in reads per million. Data is from Bhat et al., 2023 ^73^. The “SLAMseq” method distinguishes maternally deposited mRNA from mRNA transcribed *de novo* from the zygotic genome by injecting single-cell embryos with a uridine analogue which will only be incorporated in new transcripts. As can be seen from *pcna*, *polq*, and *lig3* mRNA, the levels of zygotic transcripts only rise after the maternal-to-zygotic transition around 4.5–5 hpf, but levels of all four MMEJ/fork-MMEJ factors presented here are high as early as 0.75 hpf, so are maternally deposited. RPA2 and PCNA are directly involved in fork-MMEJ but not canonical MMEJ ^41^.

**Supplementary Fig. 15.**
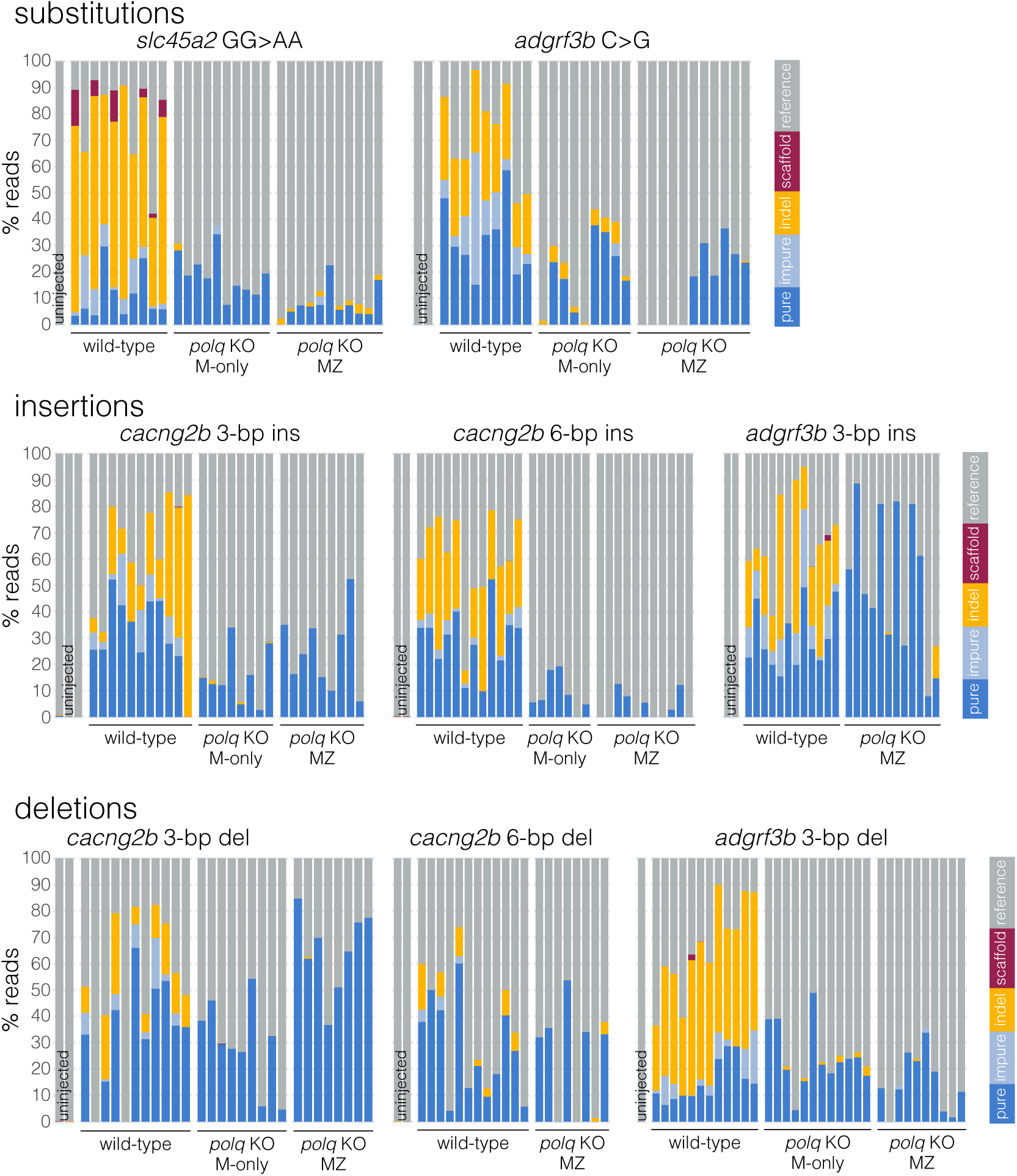
Prime editing in zebrafish embryos from homozygous *polq* KO mothers. Sequencing results from wild-type, *polq* maternal-only heterozygous KO, and *polq* maternal-zygotic homozygous KO larvae uninjected or injected with PEmax complexed with one of 8 pegRNAs (9 edits in total with *cacng2b* G>C in Fig. 4a). Each bar represents an individual larva, with colours representing the percentage of reads assigned to each label. Minimum sequencing depth was 60×. n = 224 injected samples are plotted. Wild-type *slc45a2* GG>AA samples are also plotted in Supplementary Fig. 9a. KO, knockout; M-only, maternal-only; MZ, maternal-zygotic.

**Supplementary Fig. 16.**
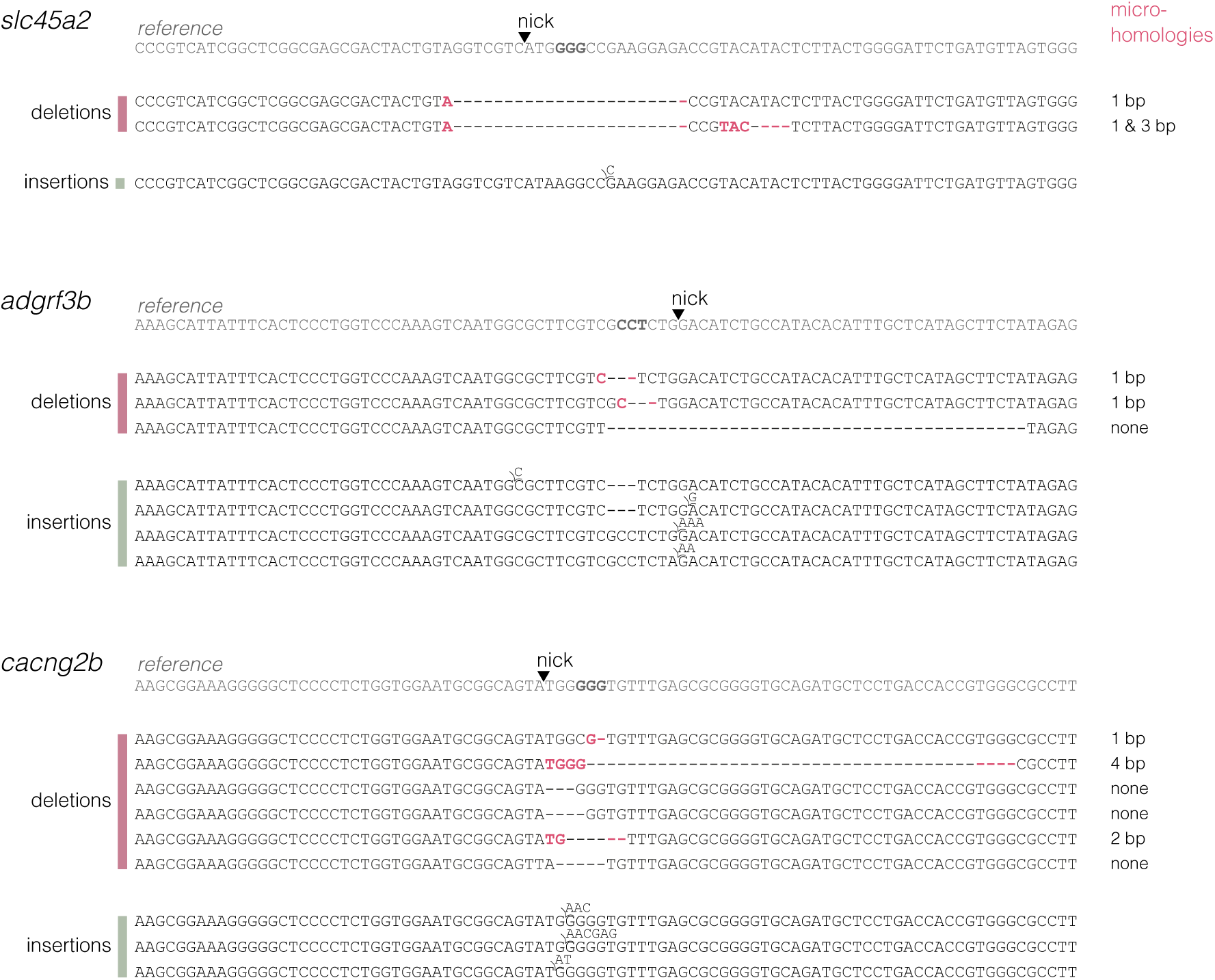
Remaining prime-editing indels in maternal-zygotic *polq* mutants. Every indel observed after prime editing in *polq* maternal-zygotic homozygous KO. For deletions, microhomologies are written in pink and right column gives their lengths. The PAM is written in bold in the reference sequences.

**Supplementary Fig. 17.**
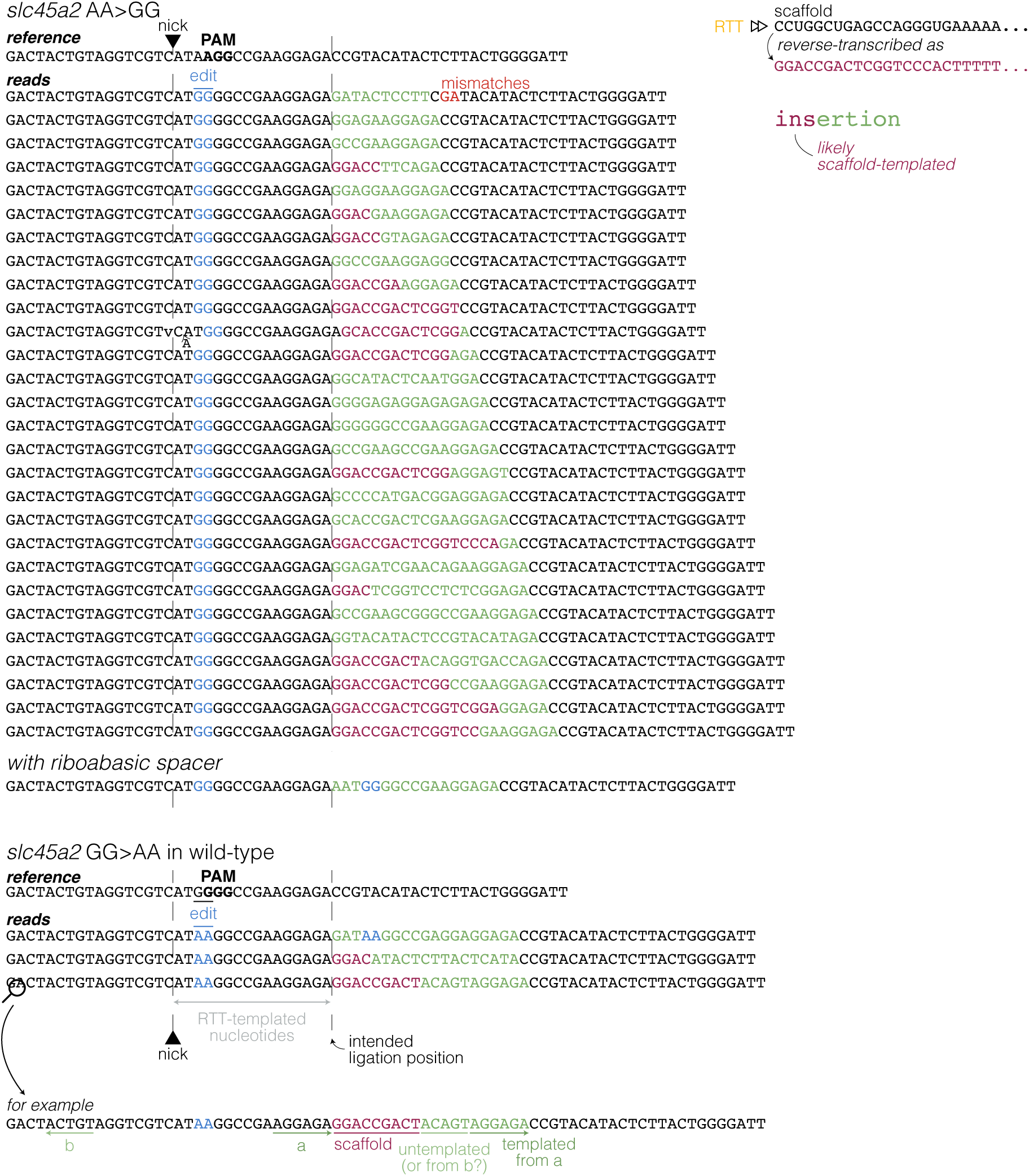
Examples of insertions at the end of the RTT-templated nucleotides. Every insertion at the *slc45a2* locus that is longer than 9 bp and starts exactly at the end of the RTT-templated nucleotides (nick + 16 bp). At the top is every unique insertion found in *slc45a2* AA>GG samples, except those injected with PEmax SpRY. At the bottom is every insertion found in *slc45a2* GG>AA samples. Inserted nucleotides are purple or green. In purple are inserted nucleotides likely templated from the scaffold because the inserted sequence starts with GGAC[…], which differentiates with GGAG[…] that could be templated from the end of the RTT-templated nucleotides.

**Supplementary Fig. 18.**
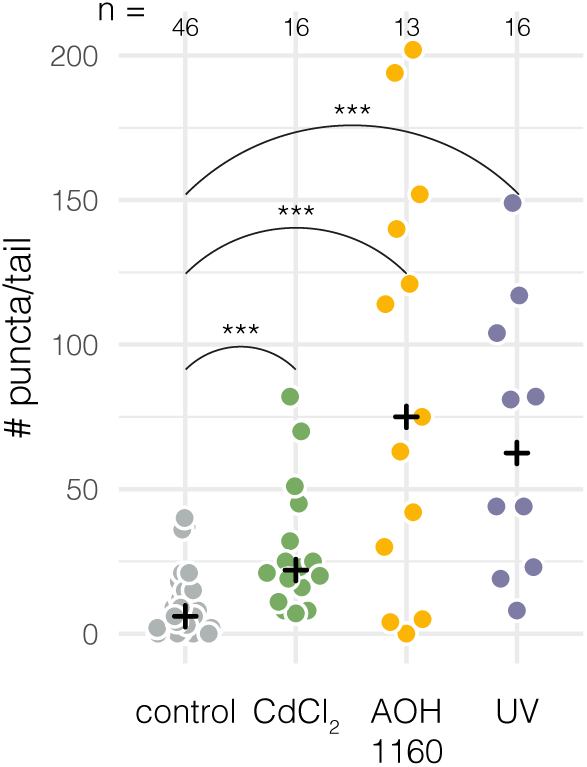
Acridine orange staining detects a rise in apoptotic cells caused by DNA damage. Detection of apoptotic cells using acridine orange staining. Each datapoint represents the number of acridine-positive puncta in the tail of an individual 1-dpf wild-type embryo, either left untreated or treated directly after fertilisation with 20 µM CdCl2 (cadmium), or 20 µM AOH1160 for 24 hr, or UV radiation (∼300 nm) for 30 min. Embryos were stained and imaged the next day at 1 dpf. Black crosses mark the group medians. *** p < 0.001 by Mann-Whitney U tests.

**Supplementary Fig. 19.**
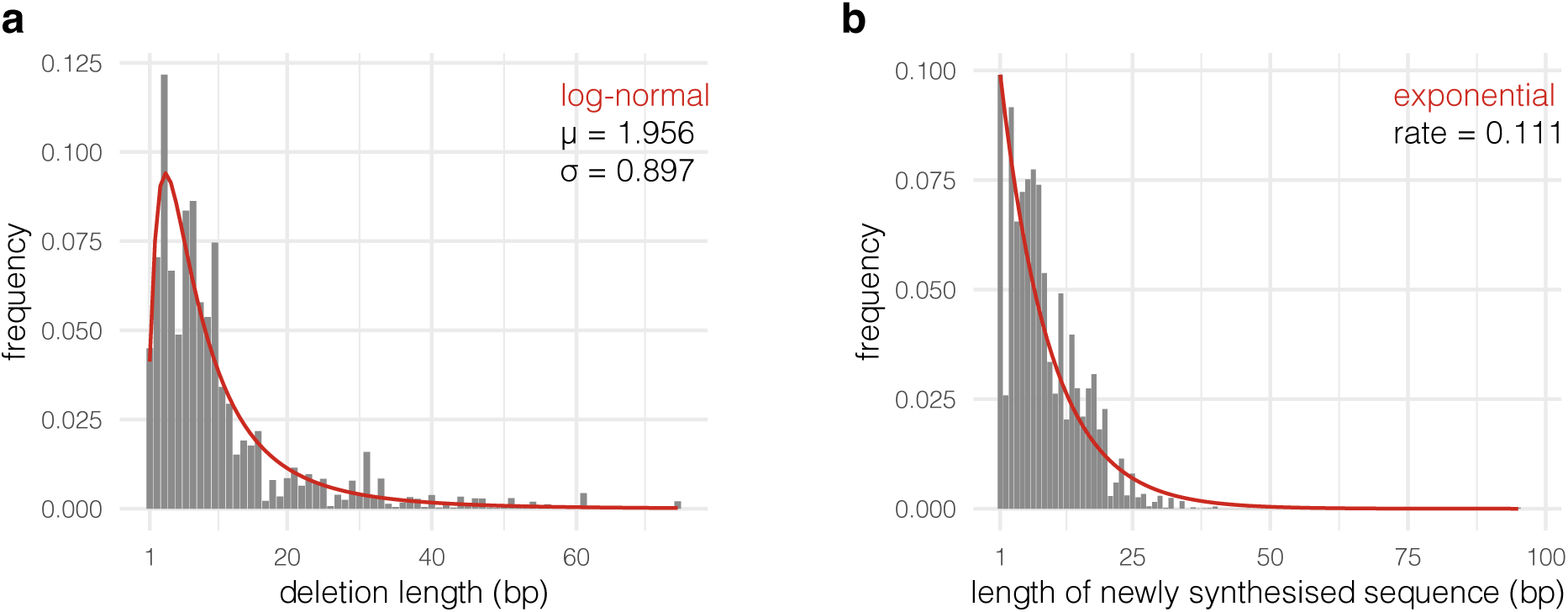
Distribution of deletion and insertion lengths after repair of Cas9 DSBs. a, Deletion length frequencies observed in a dataset of n = 191 SpCas9-injected larvae mutated at n = 38 loci. The red curve is the fitted log-normal distribution. µ, location parameter; σ, shape parameter. **b,** In zebrafish embryos, insertions generated during repair of a DSB often appear concomitantly with a multi-bp substitution (see example in Fig. 2e). We defined the “newly synthesised sequence” as the insertion together with the substitution if present. Here, frequencies of newly synthesised sequence lengths observed in a dataset of n = 191 SpCas9-injected larvae mutated at n = 38 loci. The red curve is the fitted exponential.

